# Chromosome-level genome and the identification of sex chromosomes in Uloborus diversus

**DOI:** 10.1101/2022.06.14.495972

**Authors:** Jeremiah Miller, Aleksey V Zimin, Andrew Gordus

**Affiliations:** Department of Biology, Johns Hopkins University, Baltimore, MD; Department of Biomedical Engineering, Johns Hopkins University, Baltimore, MD; Center for Computational Biology, Johns Hopkins University, Baltimore, MD; Solomon H. Snyder Department of Neuroscience, Johns Hopkins University, Baltimore, MD

## Abstract

The orb-web is a remarkable example of animal architecture that is observed in families of spiders that diverged over 200 million years ago. While several genomes exist for Araneid orb-weavers, none exist for other orb-weaving families, hampering efforts to investigate the genetic basis of this complex behavior. Here we present a chromosome-level genome assembly for the cribellate orb-weaving spider *Uloborus diversus*. The assembly reinforces evidence of an ancient arachnid genome duplication and identifies complete open reading frames for every class of spidroin gene, which encode the proteins that are the key structural components of spider silks. We identified the two X chromosomes for *U. diversus* and identify candidate sex-determining genes. This chromosome-level assembly will be a valuable resource for evolutionary research into the origins of orb-weaving, spidroin evolution, chromosomal rearrangement, and chromosomal sex-determination in spiders.

## Introduction

Spiders are among the most successful and diverse terrestrial predators on Earth. Almost 400 million years of evolution has produced more than 50,000 extant spider species representing 128 families that are distributed over every continent except Antarctica(Gloor et al. 2017). The success of these animals is due in part to the diversity of behaviors that have evolved to capture prey in different environments(Vollrath and Selden 2007). Many spiders attack their prey by physically grabbing and immobilizing them with venom and use their silk exclusively for egg sacs. Others use silk to line their burrows, or construct webs of varying geometry and composition to detect or entrap prey. The diversity of web use correlates with a diversity of spidroin proteins that form silk, as well as the glands that produce these proteins(Vollrath and Selden 2007; Blackledge et al. 2009; Gatesy et al. 2001; Foelix 2011; Vollrath 1999). Spiders such as orb-weavers alternate between glands depending upon the web feature they are constructing. For example, load bearing parts of the web such as the radii are composed of major ampullate silk that has high tensile strength, whereas the anchors are made up of pyriform silk which is sticky and amorphous(Foelix 2011).

Remarkably, the orb web is not restricted to a single monophyletic group, but is observed in two lineages that diverged 250 million years ago, leading to considerable debate about its evolutionary origins(Blackledge et al. 2009; Fernández et al. 2018; Coddington et al. 2019; Kallal et al. 2021) (**Figure 1**). Araneoidea is the largest superfamily of orb weavers, which have evolved adhesive aggregate spidroins that are used in the capture spiral to adhere prey to the web(Gatesy et al. 2001; Sahni et al. 2010; Opell and Hendricks 2010; Hayashi and Lewis 2000, 1998). However, uloborids also build orb webs, but use a more ancient cribellate spidroin in their capture spiral to immobilize prey(Peters 1992, 1984; Blackledge and Hayashi 2006; Piorkowski et al. 2020). In addition to Uloboridae, other families such as Deinopidae, and Oecobiidae + Hersiliidae (UDOH grade in **Figure 1**) also build orb-webs, but with more derived behavioral and structural characteristics(Coddington 1986). Orb-weaving is an innate behavior, with discrete stages of web construction that are shared between araneoid and non-araneoid orb-weavers(Zschokke and Vollrath 1995). When exposed to neuroactive compounds, the behaviors within specific stages are altered, indicating that the neuronal targets of these compounds are more important for certain stages than for others(Witt and Reed 1965; Hesselberg and Vollrath 2004; Reed et al. 1982). This behavioral paradigm offers an excellent model for understanding not only how complex behaviors can be organized in a small brain(Corver et al. 2021), but also how such behaviors evolve.

**Figure 1:**
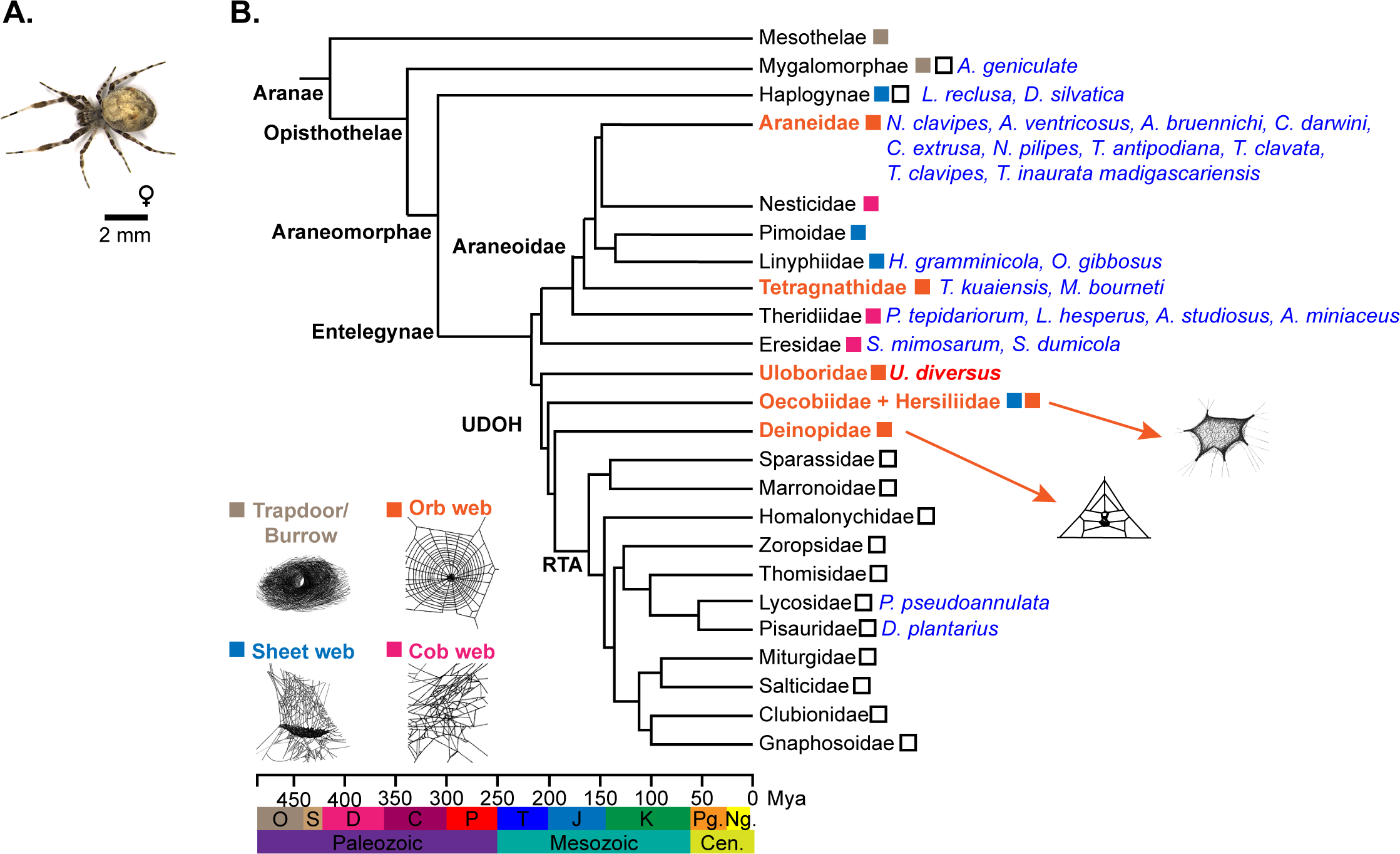
Spider Phylogeny. A) A female *U. diversus*. B) Phylogeny of spiders. Orb weaver families are highlighted in orange. Species with sequenced genomes are highlighted in blue. *U. diversus* is highlighted in red. Example webs from Rooney(Roberson et al. 2016), Glatz(Glatz 1967), Coddington(Coddington 1986). UDOH = Uloboridae, Deinopidae, Oecobiidae, Hersiliidae. RTA = Retrolateral tibial apophysis clade. O = Ordovician, S = Silurian, D = Devonian, C = Carboniferous, P = Permian, T = Triassic, J = Jurassic, K = Cretaceous, Pg. = Paleogene, Ng. = Neogene, Mya = millions of years ago

However, a genetic understanding of the evolution of orb weaving behavior is hampered by a lack of sequenced genomes for non-araneoid families. Spider genomes are enormous with high repeat content, making them challenging to assemble(Sanggaard et al. 2014; Stellwagen and Renberg 2019; Ayoub et al. 2013). While 22 spider genomes have been assembled and made publicly available(Sanggaard et al. 2014; Babb et al. 2017; Kono et al. 2019; Sheffer et al. 2021; Sánchez-Herrero et al. 2019; Schwager et al. 2017; Fan et al. 2021; Yu et al. 2019; Liu et al.2019; Escuer et al. 2022; Cerca et al. 2021; Zhu et al. 2022; Hendrickx et al. 2022; Kono et al. 2021a; Purcell and Pruitt 2019; Li et al. 2022; Kono et al. 2021b) (**Table S1**), most are highly fragmented, with 6 assembled to chromosome-scale(Sheffer et al. 2021; Fan et al. 2021; Escuer et al. 2022; Zhu et al. 2022; Hendrickx et al. 2022; Kono et al. 2021b). Of the 22 genomes, only 7 represent the Araneoidea(Kono et al. 2019; Sheffer et al. 2021; Fan et al. 2021; Kono et al. 2021a) or ecribellate orb-weavers, 2 of which have been assembled to chromosome-scale(Sheffer et al. 2021; Fan et al. 2021). There are no published genomes for the UDOH clade, and only 1 genome represents a member of the cribellate RTA clade (**Figure 1B**)(Garb et al. 2018). Chromosome-level assemblies are essential for understanding evolutionary divergence and identifying sites of chromosomal reorganization that play roles in adaptation and speciation.

Spiders have multiple sex chromosomes, with ♂X_1_X_2_/♀X_1_X_1_X_2_X_2_ being the most common sex determination observed. Sex chromosome dosage compensation has evolved multiple times, but all known genetic mechanisms are for single sex chromosome systems. A molecular understanding of dosage compensation in spiders is lacking, in part due to a paucity of sex-associated genetic loci.

*Uloborus diversus* (**Figure 1A**) of the UDOH clade, is a cribellate orb weaving spider native to the desert Southwest in the United States(Eberhard 1971) and an important model for understanding the evolution of spidroins and orb-weaving(Garb et al. 2018). The existence of a non-araneoid orb-weaving genome is crucial for addressing the evolutionary origins of orb-weaving. Recent work has demonstrated the utility of this species as a model system for understanding orb-weaving behavior(Corver et al. 2021). This, combined with the potential to compare both behaviors and their genetic underpinnings across divergent species of orb-weavers offers a rich opportunity to understand the underlying genetics that encode this behavior, and whether orb-weaving behaviors are conserved or convergently evolved. Here, we report a high-quality, chromosome-scale draft genome assembly of *Uloborus diversus* (NCBI: txid327109), as well as a complementary transcriptome assembly and gene annotations. This genome enabled the identification of full >10kb spidroin genes, as well as the identification of sex chromosomes for this species. This chromosome-level assembly will be a valuable resource for evolutionary research into the origins of orb-weaving, spidroin evolution, chromosomal rearrangement, and chromosomal sex-determination in spiders.

## Results

### Genome Sequencing

We sequenced and assembled a high-quality, chromosome-scale genome assembly for *Uloborus diversus* using a hybrid approach that leveraged the complementary benefits of multiple technologies. The genome of *U. diversus* contains long regions of low-complexity sequence, which hinders assembly using short-reads alone, as well as extremely long protein-coding genes, which makes long-reads necessary for a reference-quality assembly(Ayoub et al. 2013; Sanggaard et al. 2014; Babb et al. 2017; Kono et al. 2019; Stellwagen and Renberg 2019; Sheffer et al. 2021). Illumina sequencing provides high sequence fidelity but short read lengths, while ONT sequencing provides long read lengths, useful for scaffolding and spanning long, low complexity regions, but lower sequence fidelity(Giani et al. 2020). PacBio HiFi sequencing provides an excellent combination of long read lengths and high sequence fidelity, however, we were able to produce multiple megabase-long reads with ONT, which is not possible with PacBio. Each of these sequencing technologies provided unique advantages for improving the overall assembly.

To limit genetic variation, we used sequencing data from only 5 unmated female spiders in our assembly. We used Illumina to obtain high fidelity read data with from a single female spider, generating 795M 150 bp read pairs totaling 119.3 Gb. Because Illumina short reads are not sufficiently long to span long, highly repetitive regions encountered in spider genomes, we sequenced 3 ONT libraries, each from a single female, generating 14.7M reads totaling 98.4 Gb, with a read N50 of 6.7 kb. To obtain long sequencing reads with high sequence fidelity, we also sequenced a single adult female using PacBio HiFi, generating 35M subreads totaling 412.8 Gb with a read N50 of 12.8 kb, which yielded 2M consensus reads totaling 26 Gb with a read N50 of 13.0 kb. To investigate the sex determination system in *U. diversus*, we also generated an Illumina library from a single adult male, producing 937M 50 bp read pairs totaling 46.8 Gb. Sequencing library statistics are available in **Table 1**.

**Table 1.** Summary of Library Statistics.

| Library Type | Instrument | Mean Read Length | Number of Reads or<br>Read Pairs | Bases Sequenced | Coverage (X)<sup>a</sup> |
| --- | --- | --- | --- | --- | --- |
| Illumina | NovaSeq 6000 | 2 x 150 bp | 795 million (female) 937<br>million (male) | 119.3 Gb (female)<br>46.8 Gb (male) | 60 (female)<br>24 (male) |
| ONT | PromethION | 6.7 kb | 14.7 million | 98.4 Gb | 50 |
| PacBio | Sequel II | 12.8 kb (subreads)<br>13.0 kb (consensus) | 34.9 million (subreads)<br>2.0 million (consensus) | 412.8 Gb (subreads)<br>26.2 Gb (consensus) | 208 (subreads)<br>13.1 (consensus) |
| Chicago Hi-C | HiSeq X | 2 x 150 bp | 286 million | 85.8 Gb | 21b<br>43c |
| Dovetail Hi-C | HiSeq X | 2 x 150 bp | 537 million | 161.1 Gb | 1,009b<br>81c |

<sup>a</sup> Based on in silico genome size estimate of 1.98 Gb by *k*-mer analysis with GenomeScope 2.0
<sup>b</sup> Physical coverage, defined as the number of read pairs that span a base pair
<sup>c</sup> Sequence coverage, defined as number of times a base pair is directly observed in sequencing data

### Genome Size, Heterozygosity, and Coverage Estimation

To assess the size and heterozygosity of the genome, we used Jellyfish(Marçais and Kingsford 2011) to count the frequency of canonical 21-mers in our adult female Illumina sequencing reads and used the 21-mer distribution as input to GenomeScope(Ranallo-Benavidez et al. 2020). The resulting model estimated a genome size of 1.98 Gb with a heterozygosity of 1.38% and 50.2% of the genome occurring as unique sequence (**Figure 2**), similar to other spider genomes (**Supplementary Table S1**)(Sanggaard et al. 2014; Babb et al. 2017; Kono et al. 2019; Sheffer et al. 2021; Sánchez-Herrero et al. 2019; Schwager et al. 2017; Fan et al. 2021; Yu et al. 2019; Liu et al. 2019). Given this genome size, our Illumina sequencing yielded 63x coverage, our ONT sequencing yielded 68x coverage, and our PacBio HiFi yielded 208x in raw read coverage and 13x in consensus read coverage (**Table 1**).

**Figure 2:**
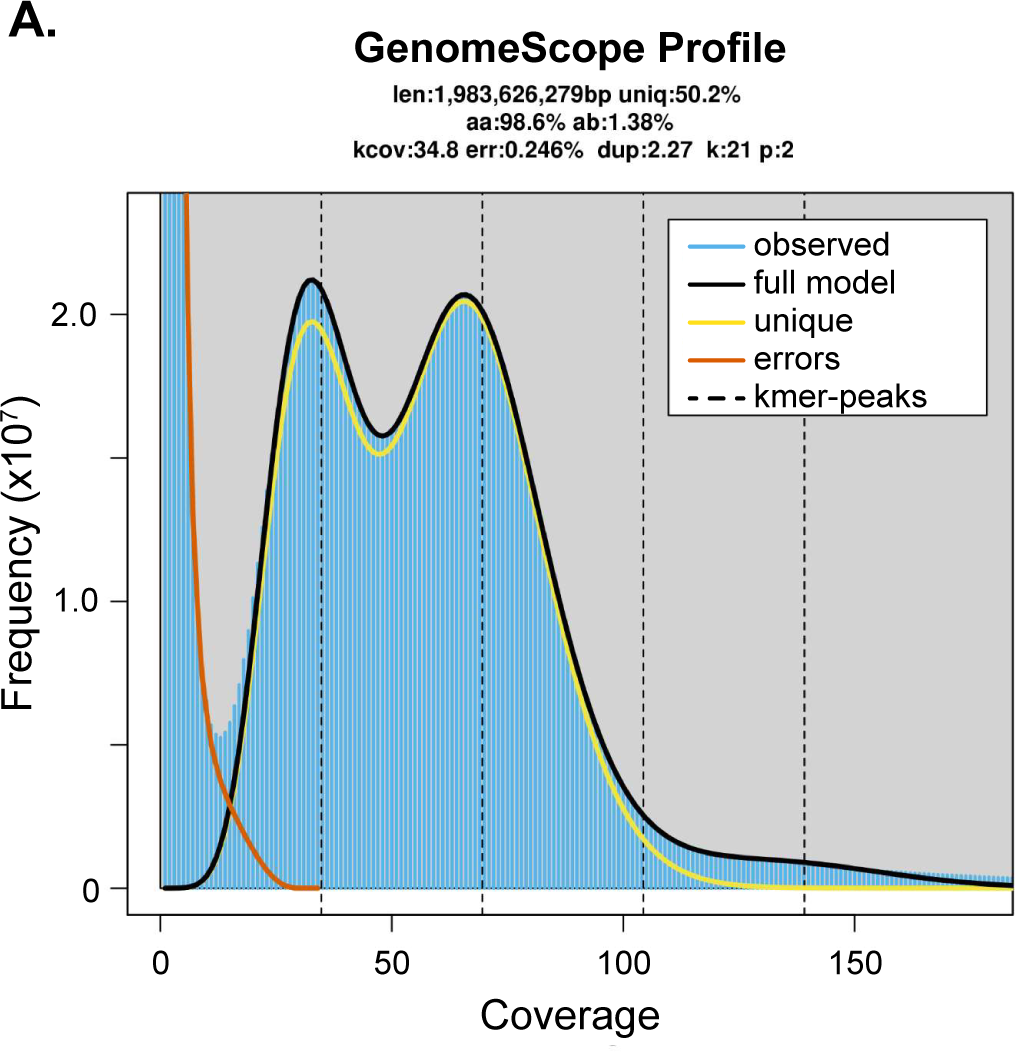
GenomeScope Plot from Illumina Data. A) Kmer spectra for Illumina reads from a single, virgin female. The diploid and haploid peaks are at 70x and 35x coverage, respectively.

### Karyotype of *U. diversus*

To infer the expected number of pseudo-chromosomes in our final assembly, we determined the number of chromosomes in *U. diversus* using metaphase karyotyping. Mitotic chromosome spreads from developing embryos displayed two distinct patterns of chromosome number; either 18 or 20 (**Figure 3A**), consistent with ♂X_1_X_2_/♀X_1_X_1_X_2_X_2_ sex determination, which is the most common form of sex determination observed in spiders(Sember et al. 2020). Thus, *U. diversus* appears to have 8 autosomes and 2 sex chromosomes.

**Figure 3:**
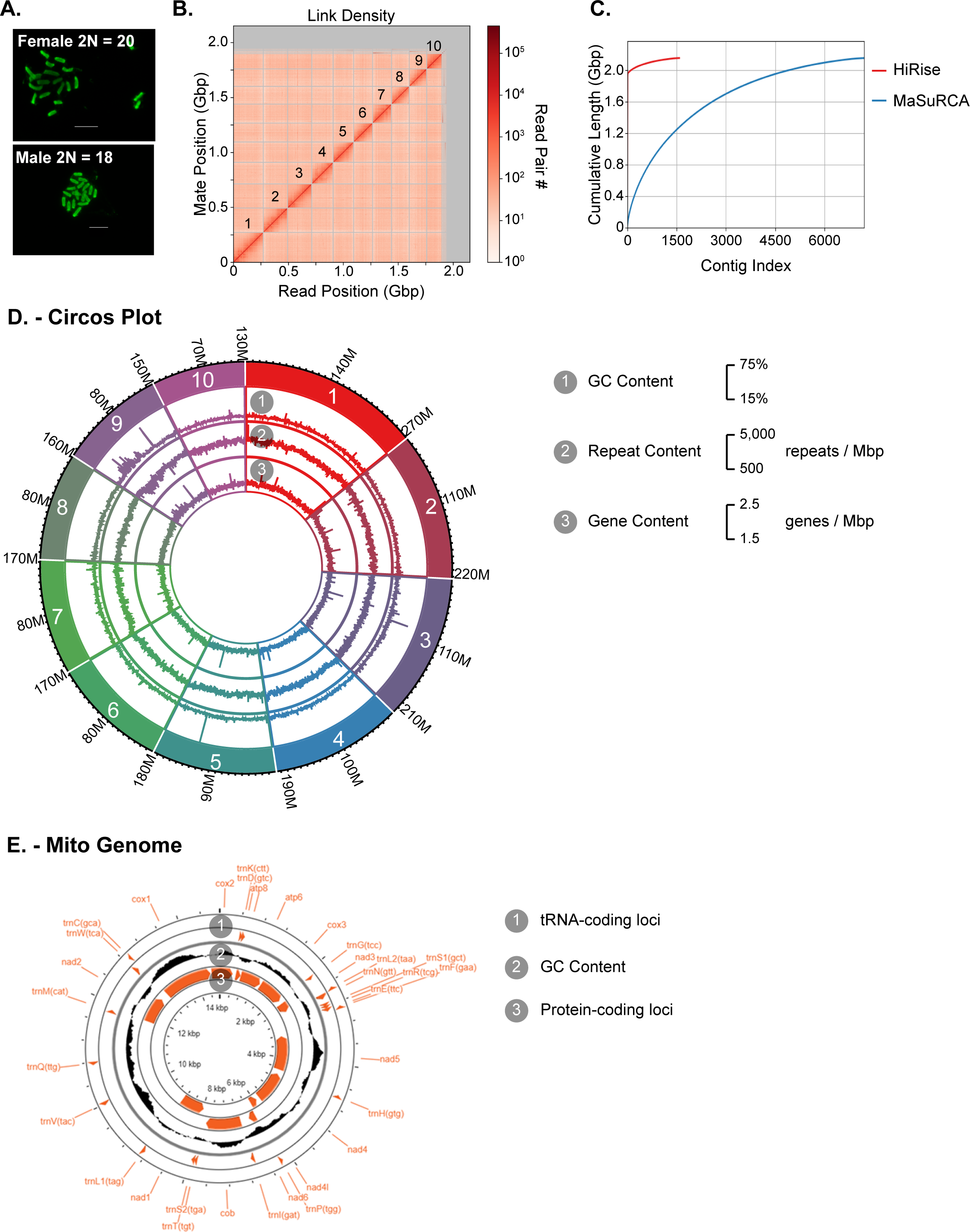
Chromosome Scale Genome Assembly. A) Karyotype of female and male embryos. Female and male diploid sizes of 20 and 18, respectively, indicate a ♂X1X2/♀X1X1X2X2 sex-determination system, with 8 autosomes. B) Hi-C linkage map of assembled scaffolds. The 10 largest scaffolds are annotated. C) Comparison of HiRise and MaSuRCA assemblies. The majority of the HiRise assembly is captured by the first 10 scaffolds. D) Circos plot of 10 largest nuclear scaffolds, highlighting GC content, repeat content, and gene content across the scaffolds. E) Circos plot of mitochondrial scaffold, highlighting tRNA-coding loci, protein-coding loci, and GC content.

### *De novo* Nuclear Genome Assembly

First, we used MaSuRCA(Zimin et al. 2013, 2017) to produce an initial assembly (*U. diversus* v.1.0) using Illumina short-read data scaffolded by ONT long-read data, consisting of 68,259 scaffolds spanning 3.22 Gb. The scaffold N50 was 98,014 bp and the scaffold L50 was 6,558 (**Table 2**), with 94.7% of complete BUSCOs (**Table 3**). The inferred redundancy accounts for the significant increase in the length of the assembly compared to the expected genome size. High heterozygosity leads to alternative haplotypes that can often be misassembled into their own contigs.

**Table 2.** Summary of Genome Assembly Statistics.

|  | *U. div v.1.0* |  | *U. div v.1.1* |  | *U. div v.1.2* |  | *U. div v.1.3* |  | *U. div v.2.0* |  | *U. div v.3.0* |  | *U. div v.3.1* |  |
| --- | --- | --- | --- | --- | --- | --- | --- | --- | --- | --- | --- | --- | --- | --- |
|  | Contigs | Scaffolds | Contigs | Scaffolds | Contigs | Scaffolds | Contigs | Scaffolds | Contigs | Scaffolds | Contigs | Scaffolds | Contigs | Scaffolds |
| Total Length | 3,218,774,062 |  | 3,218,805,127 |  | 2,798,889,957 |  | 2,802,061,857 |  | 2,122,354,655 |  | 2,151,304,433 |  | 2,151,890,133 |  |
| Number of Contigs / Scaffolds | 68,581 | 68,259 | 68,581 | 66,417 | 50,738 | 48,632 | 55,142 | 21,317 | 9,997 | 9,734 | 7,467 | 7,197 | 7,713 | 1,586 |
| Largest Contig / Scaffold | 3,987,394 | 3,987,394 | 3,987,394 | 6,446,995 | 3,987,394 | 6,446,995 | 3,987,394 | 367,325,988 | 4,617,239 | 4,617,239 | 5,877,357 | 5,877,357 | 5,877,357 | 272,330,431 |
| Mean Contig / Scaffold Length | 46,933 | 47,155 | 46,933 | 48,463 | 55,162 | 57,552 | 50,756 | 131,447 | 212,298 | 218,035 | 288,107 | 298,916 | 278,918 | 1,356,803 |
| Median Contig / Scaffold Length | 21,841 | 21,923 | 21,841 | 21,546 | 23,184 | 22,648 | 24,516 | 12,994 | 137,429 | 218,035 | 166,676 | 175,879 | 168,537 | 94,191 |
| Smallest Contig / Scaffold | 1,106 | 1,667 | 1,106 | 1,667 | 1,106 | 1,667 | 10 | 312 | 10,045 | 10,045 | 10,045 | 10,045 | 380 | 10,220 |
| N10 / L10 | 558,595 / 374 | 564,850 / 373 | 558,595 / 374 | 686,213 / 308 | 613,610 / 302 | 744,166 / 249 | 432,729 / 4,437 | 367,325,988 / 1 | 1,100,324 / 132 | 1,100,324 / 132 | 1,705,719 / 86 | 1,715,742 / 84 | 1,564,267 / 91 | 272,330,431 / 1 |
| N20 / L 20 | 330,632 / 1,135 | 332,714 / 1,131 | 330,632 / 1,135 | 395,570 / 943 | 375,534 / 894 | 452,633 / 744 | 271,565 / 1,278 | 289,477,137 / 2 | 711,683 / 383 | 713,631 / 382 | 1,147,222 / 244 | 1,163,865 / 241 | 1,032,642 / 237 | 221,850,441 / 2 |
| N30 / L30 | 214,753 / 2,358 | 215,989 / 2,348 | 214,753 / 2,358 | 252,183 / 1,983 | 252,860 / 1,816 | 296,524 / 1,521 | 192,451 / 2,518 | 264,078,423 / 3 | 516,155 / 741 | 518,859 / 736 | 826,976 / 467 | 844,177 / 461 | 763,983 / 509 | 218,885,758 / 3 |
| N40 / L40 | 114,360 / 4,189 | 145,056 / 4,171 | 114,360 / 4,189 | 166,010 / 3,572 | 178,542 / 3,143 | 205,473 / 2,661 | 139,664 / 4,234 | 243,450,885 / 4 | 401,815 / 1,210 | 404,285 / 1,205 | 636,610 / 763 | 648,167 / 753 | 587,015 / 831 | 185,519,777 / 4 |
| N50 / L50 | 97,524 / 6,913 | 98,015 / 6,885 | 97,524 / 6,913 | 108,431 / 5,994 | 126,236 / 5,019 | 142,390 / 4,302 | 103,046 / 6,850 | 241,033,576 / 5 | 326,127 / 1,798 | 328,082 / 1,789 | 487,746 / 1,150 | 496,769 / 1,133 | 452,781 / 1,250 | 185,519,777 / 5 |
| N60 / L60 | 64,898 / 10,992 | 65,255 / 10,942 | 64,898 / 10,992 | 69,185 / 9,724 | 88,433 / 7,678 | 97,899 / 6,679 | 75,059 / 9,775 | 217,901,867 / 7 | 259,595 / 2,527 | 261,845 / 2,511 | 380,517 / 1,651 | 387,963 / 1,626 | 359,368 / 1,784 | 172,099,698 / 7 |
| N70 / L70 | 42,949 / 17,131 | 43,310 / 17,041 | 42,949 / 17,131 | 44,720 / 15,580 | 60,079 / 11,529 | 64,377 / 10,226 | 53,001 / 14,221 | 213,666,783 / 8 | 201,138 / 3,460 | 204,029 / 3,433 | 292,644 / 2,296 | 298,089 / 2,258 | 280,097 / 2,463 | 161,310,338 / 8 |
| N80 / L80 | 28,594 / 26,353 | 28,760 / 26,208 | 28,594 / 26,353 | 29,173 / 24,561 | 38,159 / 17,400 | 39,689 / 15,782 | 35,344 / 20,692 | 183,225,947 / 9 | 148,105 / 4,686 | 151,467 / 4,638 | 206,980 / 3,169 | 211,349 / 3,116 | 200,858 / 3,369 | 159,483,530 / 9 |
| N90 / L90 | 18,127 / 40,511 | 18,198 / 40,294 | 18,127 / 40,511 | 18,295 / 38,527 | 21,027 / 27,290 | 21,374 / 25,391 | 20,312 / 31,107 | 37,840 / 1,399 | 97,979 / 6,436 | 100,002 / 6,354 | 127,987 / 4,484 | 131,815 / 4,398 | 126,330 / 4,713 | 9,355,840 / 14 |
| N100 / L100 | 1,106 / 68,581 | 1,667 / 68,259 | 1,106 / 68,581 | 1,667 / 66,417 | 1,106 / 50,738 | 1,667 / 48,632 | 10 / 55142 | 312 / 21,317 | 10,045 / 9,997 | 10,045 / 9,734 | 10,045 / 7,467 | 10,045 / 7,197 | 380 / 7,713 | 10,220 / 1,586 |
| Gaps | 322 |  | 2,164 |  | 2,106 |  | 33,825 |  | 263 |  | 270 |  | 6,127 |  |
| Ns | 32,200 |  | 63,265 |  | 57,963 |  | 3,229,863 |  | 6,049 |  | 6,749 |  | 592,449 |  |
| GC Content (%) | 33.78 |  | 33.78 |  | 33.72 |  | 33.72 |  | 33.83 |  | 33.82 |  | 33.82 |  |

U. div v.1.0 is the MaSuRCA assembly U. div v.1.1 is the MaSuRCA assembly with further scaffolding using Rascaf U. div v.1.2 is the MaSuRCA assembly with Rascaf and with reduction of redundancy using Pseudohaploid U. div v.1.3 is the MaSuRCA assembly with Rascaf and Pseudohaploid further scaffolded using Chicago and Dovetail Hi-C U. div v.2.0 is the PacBio IPA assembly U. div v.3.0 is the MaSuRCA assembly and PacBio IPA assembly merged using MaSuRCA U. div v.3.1 is the merged MaSuRCA/IPA assembly further scaffolded using Dovetail Hi-C

**Table 3.** Summary of Genome Assembly BUSCO Scores.

|  | *U. div v.1.0* | *U. div v.1.1* | *U. div v.1.2* | *U. div v.1.3* | *U. div v.2.0* | *U. div v.3.0* | *U. div v.3.1* |
| --- | --- | --- | --- | --- | --- | --- | --- |
| Complete | 94.7 | 95.4 | 95.1 | 97.0 | 92.4 | 94.1 | 94.8 |
| Single Copy | 70.9 | 72.9 | 78.4 | 84.6 | 82.1 | 82.9 | 85.2 |
| Duplicated | 23.8 | 22.5 | 16.7 | 12.4 | 10.3 | 11.2 | 9.6 |
| Fragmented | 2.3 | 1.9 | 2.0 | 1.2 | 1.7 | 1.3 | 1.0 |
| Missing | 3.0 | 2.7 | 2.9 | 1.8 | 5.9 | 4.6 | 4.2 |

Total BUSCOs 2,934
U. div v.1.0 is the MaSuRCA assembly
U. div v.1.1 is the MaSuRCA assembly with further scaffolding using Rascaf
U. div v.1.2 is the MaSuRCA assembly with Rascaf and with reduction of redundancy using Pseudohaploid
U. div v.1.3 is the MaSuRCA assembly with Rascaf and Pseudohaploid further scaffolded using Chicago and Dovetail Hi-C
U. div v.2.0 is the PacBio IPA assembly
U. div v.3.0 is the MaSuRCA assembly and PacBio IPA assembly merged using MaSuRCA
U. div v.3.1 is the merged MaSuRCA/IPA assembly further scaffolded using Dovetail Hi-C

Next, we used Rascaf(Song et al. 2016) to improve continuity and ordering of scaffolds in the initial MaSuRCA assembly. Rascaf uses paired-end RNA-seq reads to improve the contiguity of gene models and scaffolds. We observed a modest improvement, reducing the number of scaffolds from 68,259 to 63,265 with the scaffold N50 increasing from 98,014 bp to 108,431 bp and the scaffold L50 decreasing from 6,885 to 5,994, with no change in the assembly span (**Table 2**). However, despite identifying 95.4% of BUSCOs, 22.5% were duplicated (**Table 3**). This, combined with the large span of the genome, indicated a high degree of redundancy in the assembly.

To filter out redundant heterozygous contigs we used Pseudohaploid(Chen et al. 2019b). Pseudohaploid filters suspected homologous contigs and selects a single representative contig where high rates of heterozygosity prevent assemblers from appropriately identifying haplotypes. We reduced the number of scaffolds by 23.1%, with an increase in scaffold N50 and a decreased span from 3.22 Gb to 2.8 Gb (**Table 2**) and a drop in duplicated BUSCOs to 16.7% (**Table 3**). This suggests that Pseudohaploid was able to accurately collapse much of the redundant sequence attributable to alternative haplotypes.

To further improve our assembly, we sequenced using PacBio HiFi and assembled the resulting HiFi reads with PacBio’s IPA (Improved Phased Assembly) pipeline, resulting in a substantial decrease in the new assembly span, from 2.8 Gbp to only 2.1 Gbp. The number of scaffolds in this assembly was only 9,734, a remarkable improvement. The scaffold N50 in the IPA assembly was 328,082 bp and the scaffold L50 in the IPA assembly was 1,789 (**Table 2**). The BUSCO score for the IPA assembly indicated that this assembly contained 92.3% of BUSCOs complete (83.7% in single copy, 9.3% duplicated) (**Table 3**).

We then used SAMBA(Zimin and Salzberg 2022) to merge the previous MaSuRCA assembly with the IPA assembly. SAMBA used scaffolds from the MaSuRCA assembly to merge the scaffolds in the IPA assembly. The new assembly had a length of 2.1 Gbp from only 7,197 scaffolds, with a scaffold N50 of 496,769 bp (**Table 2**), and 94% complete BUSCOs (**Table 3**).

To generate a chromosome-level assembly, we used HiRise to scaffold the IPA+MaSuRCA assembly with a Dovetail Hi-C library(Putnam et al. 2016). Scaffolding did not change the amount of sequencing in the assembly; however, the total number of scaffolds was reduced by 78% to only 1,586 scaffolds, with a remarkable improvement in scaffold N50, which increased to 185,519,777 bp in the Hi-C scaffolded assembly (**Table 2**, **Figure 3C**). Most importantly, 88% of the total assembly was represented by 10 large scaffolds that comprise 1.9 Gbp (**Figure 3B**), matching the expected number of chromosomes (**Figure 3A**). The BUSCO score for the final assembly showed 94.8% of the BUSCOs were complete (with 85.2% in single copy, 9.6% duplicated) (**Table 3**). Our final chromosome-level genome assembly statistics are consistent with previously published spider genomes, with 10 scaffolds representing the 10 chromosomes and high scaffold N50(Sanggaard et al. 2014; Babb et al. 2017; Kono et al. 2019; Sheffer et al. 2021; Sánchez-Herrero et al. 2019; Schwager et al. 2017; Fan et al. 2021; Yu et al. 2019; Liu et al. 2019) (**Supplementary Table S1**). We will refer to these 10 scaffolds as pseudochromosomes.

### Repeat Annotation

To characterize repetitive sequences, we constructed a species-specific repeat library using RepeatModeler2(Flynn et al. 2020) This library was used in conjunction with the RepBase RepeatMasker Edition(Bao et al. 2015) database for masking the genome. RepeatMasker analysis of the combined *U. diversus* and RepBase repeats masked 66.6% of the final *U. diversus* genome assembly. Many (29.27%) of the repetitive regions were unclassified; however, DNA transposons accounted for a similar proportion (22.73%). Retroelements accounted for a much smaller proportion (7.7%). Total interspersed repeats account for 59.7% and simple repeats cover 2.61% of the genome (**Table 4**). The disparity between the GenomeScope estimation of 49.8% repetitive sequence and the RepeatMasker estimation of 66.6% suggests that the repeat content may be underestimated by GenomeScope. Therefore, the genome size may also be underestimated by GenomeScope. Because the length of the *U. diversus* genome is slightly less than the prediction by GenomeScope, this possibility suggests that the length of the assembly may more closely represent the true length of the *U. diversus* genome than the GenomeScope prediction. The repeat content is typical of spider genomes (**Supplemental Table S2**).

**Table 4.** Summary of Repeat Content.

| Type of Element | Number of Elements | Total Length | Percent of Assembly |
| --- | --- | --- | --- |
| Retroelements | 311,947 | 165,713,554 | 7.70 |
| SINEs | 87,037 | 46,791,087 | 2.17 |
| Penelope | 29,911 | 11,105,124 | 0.52 |
| LINEs | 151,953 | 66,189,386 | 3.08 |
| CRE/SLAC | 0 | 0 | 0.00 |
| L2 / CR1 / Rex | 31,294 | 18,030,662 | 0.84 |
| R1 / LOA / Jockey | 17,662 | 9,587,620 | 0.45 |
| R2 / R4 / NeSL | 0 | 0 | 0.00 |
| RTE / Bov-B | 40,080 | 15,062,698 | 0.70 |
| L1 / CIN4 | 21,465 | 5,682,132 | 0.26 |
| LTR elements | 72,957 | 52,733,081 | 2.45 |
| BEL / Pao | 10,210 | 8,094,671 | 0.38 |
| Ty1 / Copia | 24,043 | 10,565,700 | 0.49 |
| Gypsy / DIRS1 | 27,837 | 29,214,338 | 1.36 |
| Retroviral | 10,867 | 4,858,372 | 0.23 |
| DNA transposons | 1,510,089 | 489,113,225 | 22.73 |
| hobo-Activator | 545,025 | 164,842,101 | 7.66 |
| Tc1-IS630-Pogo | 428,233 | 150,644,883 | 7.00 |
| En-Spm | 0 | 0 | 0.00 |
| MuDR-IS905 | 0 | 0 | 0.00 |
| PiggyBac | 11,411 | 4,281,311 | 0.20 |
| Tourist / Harbinger | 6,573 | 2,466,806 | 0.11 |
| Other (Mirage, P-element, Transib) | 2,942 | 1,822,173 | 0.08 |
| Rolling-circles | 229,864 | 88,460,474 | 4.11 |
| Unclassified | 2,482,847 | 629,886,344 | 29.27 |
| Total Interspersed Repeats |  | 1,284,713,123 | 59.70 |
| Small RNA | 20,797 | 5,420,377 | 0.25 |
| Satellites | 0 | 0 | 0.00 |
| Simple repeats | 529,329 | 56,254,893 | 2.61 |
| Low complexity | 67,980 | 3,208,338 | 0.15 |
| **Total:** |  |  | **66.58** |

### Transcriptome Sequencing and Assembly

To identify protein-coding genes, we assembled a transcriptome. To capture a wide range of transcripts, we extracted RNA from spiders at multiple developmental stages and from male and female adults. We produced an Illumina short-read sequencing library from: a whole adult female, a whole adult male, the dissected prosoma (cephalothorax) from an adult female, the dissected opisthosoma (abdomen) from the same adult female, the dissected prosoma from an adult male, the dissected opisthosoma from the same adult male, the pooled legs from the dissected male and female, a single 4th instar female, and approximately 30 pooled 2nd instars. 302M read pairs were generated, totaling 45.4 Gbp. We used Trinity(Haas et al. 2013) to assemble a genome-guided transcriptome. We then used TransDecoder(Haas et al. 2013) to find coding regions within our transcripts. We included homology searches to known proteins using both BLAST (Basic Local Alignment Search Tool)(Altschul et al. 1990; Altschul 1997) and Pfam(Mistry et al. 2021) searches. We assessed the BUSCO score of the long ORFs predicted by TransDecoder, finding that 90.9% of the BUSCOs were present and complete, with 54.8% single copy and 36.1% duplicated, with 1.5% present but fragmented and 7.6% missing. (**Table 3**).

### Protein-Coding Gene Annotation

For protein coding gene annotations, we used BRAKER2(Hoff et al. 2016; Brůna et al. 2021) with our RNAseq data and homology evidence using a custom library of spider proteins obtained from NCBI (**Supplemental Table S3**). The number of predicted genes in the final *U. diversus* assembly was 44,408; with 40,466 models predicted on the 10 pseudochromosomes (**Table 5**), with 86.7% of complete BUSCOs. To functionally annotate these genes, we used Interproscan(Quevillon et al. 2005; Jones et al. 2014) to annotate the longest CDS for each gene. 30,911 models were assigned a domain or function from one or more of the databases used (**Table 6**).

**Table 5.** Summary of Annotation Statistics.

|  |  |
| --- | --- |
| Number of Gene Models | 45,762 |
| Minimum Gene Model Length (bp) | 60 |
| Maximum Gene Model Length (bp) | 408,348 |
| Average Gene Model Length (bp) | 16,764 |
| Number of Exons | 222,483 |
| Average Number of Exons per Gene Model | 5 |
| Average Exon Length (bp) | 237 |
| Number of Transcripts | 47,540 |
| Average Number of Transcripts per Gene Model | 1 |
| Number of Gene Models < 200 bp | 37 |

**Table 6.** Summary of InterproScan Results.

| Database | Total Hits | Individual mRNAs with Hits |
| --- | --- | --- |
| CDD | 11047 | 6560 |
| Coils | 9254 | 5947 |
| Gene3D | 34377 | 16424 |
| Hamap | 270 | 260 |
| MobiDBLite | 44028 | 14046 |
| PANTHER | 46497 | 20941 |
| Pfam | 35843 | 19491 |
| PIRSF | 916 | 736 |
| PRINTS | 16380 | 3134 |
| ProSitePatterns | 8222 | 3898 |
| ProSiteProfiles | 23252 | 9922 |
| SFLD | 120 | 62 |
| SMART | 25638 | 7561 |
| SUPERFAMILY | 26766 | 15559 |
| TIGRFAM | 821 | 736 |

### Non-Coding RNA Annotation

We used tRNAscan-SE(Chan and Lowe 2019; Chan et al. 2021) to annotate transfer RNAs. We found 3,084 tRNAs coding for the standard 20 amino acids and 14 tRNAs coding for selenocysteine (TCA) tRNAs. We found 21 tRNAs with undetermined or unknown isotypes, 537 tRNAs with mismatched isotypes, and 57,824 putative tRNA pseudogenes. We identified no putative suppressor tRNAs. We used Barrnap(Seeman 2018) to annotate ribosomal rRNAs. We found 114 rRNAs, of which 100 were located on the 10 pseudo-chromosomes. These included: 6 copies of the 18S subunit, with 4 on pseudochromosomes; 83 copies of the 5S subunit, with 81 on pseudochromosomes; 6 copies of the 5.8S subunit, with 2 on pseudochromosomes; and 19 copies of the 28S subunit, with 13 pseudochromosomes.

### Mitogenome Assembly

Animal mitochondrial genomes comprise 37 genes: 13 protein coding genes, 22 tRNAs, 2 rRNAs, and at least one control region(Boore 1999). We assembled the mitochondrial genome sequence with NOVOplasty(Dierckxsens et al. 2016) using the adult female Illumina DNA seq data and each of the mitochondrial genome sequences listed in **Supplemental Table S4** as sources for seed sequences(Wang et al. 2019b; Zhu and Zhang 2017; Wang et al. 2014; Liu et al. 2015; Fang et al. 2016; Masta and Boore 2008; Kumar et al. 2020; Li et al. 2016; Qiu et al. 2005; Pan et al. 2014; Kim et al. 2014; Pan et al. 2016; Tian et al. 2016; Wang et al. 2016a). Each run produced the same single, circularized 14,737 bp mitochondrial sequence, consistent with the expected size for an arachnid mitochondrial sequence(Boore 1999). We annotated the sequence with the MITOS2 web server(Bernt et al. 2013) and found all 13 of the expected protein coding genes, 20 of 22 tRNAs, 2 rRNAs, and the control region (**Fig. 3E**). All identified tRNAs were truncated and lacked T-arms, which is unique to spiders and has been observed in other species(Masta and Boore 2008; Wang et al. 2016b; Li et al. 2016; Pons et al. 2019).

### Identification and Analysis of Spidroins

Spidroins are a unique class of proteins that are the primary components of spider silk. While all spiders produce silk, spidroins have evolved for different uses in web-making. Orb-weavers in particular evolved several silk glands that each produce a different repertoire of spidroins to make different silks with varying utility. Several ecribellate Araneidae spidroins have been sequenced, and many of these spidroins are also made by cribellate orb-weavers such as *U. diversus*. However, ecribellate spiders evolved a unique type of hydrated flageliform silk for their capture spiral, whereas cribellate spiders such as *U. diversus* use a dry cribellate silk in their capture spirals.

Spider dragline silk has the strongest stress and strain capabilities of any known substance. Interest in silk properties extends beyond their evolved use, as silk has many potential human applications in both industry and medicine(Kumari et al. 2020; Xu et al. 2019; Öksüz et al. 2021; Choi and Choy 2020; Mayank et al. 2022; Liu et al. 2020; Lewis 2006; Teulé et al. 2007). However, the genetic characterization of spidroins is often challenging due to their exceptional length (coding regions >5 kb) and high repeat content(Stellwagen and Renberg 2019). The annotation of these genes is difficult and often fragmented because reads rarely span the entire length of these genes. With the exceptional contiguity and read depth of our assembly, due to the diversity of sequencing technologies employed, we identified the entire open-read frames of all major spidroins in the *U. diversus* genome.

We found 10 full-length candidate sequences, including at least one candidate for each of the seven types of spidroin used by cribellate orb-weavers. There were no gaps in the assembly interrupting any of our candidate spidroin sequences. We performed read mapping to validate the continuity of each full-length sequence and ensure that the predicted sequences were not chimeric. In 8 cases, including 3 minor spidroin (MiSp) candidates, a major spidroin (MaSp) MaSp-1 candidate, 2 MaSp-2 candidates, a tubuliform spidroin (TuSp) candidate, and an aciniform spidroin (AcSp) candidate, the full length of the predicted genomic region was spanned entirely by at least one HiFi consensus read. In the remaining 3 cases, consisting of a pyriform spidroin (PySp) candidate, a cribellate spidroin CrSp candidate, and a pseudoflagelliform spidroin (Pflag) candidate, no more than 2 HiFi reads were necessary to span the entirety of the predicted genomic region, and in each of these cases there was sufficient depth and overlap in the reads to call the region with high confidence (see **Supplemental Figure S1**). The length of the coding regions for the spidroins ranged from 5.5 kb to 20 kb. This is consistent with expectations of full-length sequences found in other spiders(Babb et al. 2017; Kono et al. 2019, 2020). Only in the case of MaSp-1 were we unable to call a complete, full-length sequence. The finalized spidroin annotations included only two spidroins with multiple exons: CrSp and MaSp-1; however, the structure of MaSp1 remains unclear. All other spidroins were found to be single exon sequences. While most single-exon genes tend to be small highly expressed proteins such as histones, spidroins are a rare exception. The single exon structure of spidroin genes has been noted in other species(Ayoub et al. 2013; Kono et al. 2020; Garb and Hayashi 2005; Motriuk-Smith et al. 2005; Ayoub and Hayashi 2008; Liu et al. 2022; Wen et al. 2020; Wang et al. 2019a; Wen et al. 2017).

#### Aciniform spidroin (AcSp)

Aciniform silk is one of the toughest spider silks, and is typically used for wrapping prey(Tremblay et al. 2015). A single exon for AcSp was identified on Chromosome 7 (**Figure 4A** and **Table 7**), consistent with the sequences in the Araneid orb-weaving spiders *Araneus ventricosus*(Wen et al. 2018) and *Argiope agentata*(Chaw et al. 2014), as well as the cobweb spider *Latrodectus hesperus*(Ayoub et al. 2013). Our confidence in this sequence is high since the complete sequence was spanned entirely by multiple PacBio HiFi reads. In the repetitive region, we found 13 iterated repeats of a 357 amino acid motif, and a 14th partial repeat (**Supplemental Figure S1**). As with previous reports on the structure of AcSp(Ayoub et al. 2013; Hayashi 2004; Chaw et al. 2014; Wen et al. 2018), we also found that the repeats are remarkably well-homogenized. After removal of the signal peptide between Ser-23 and Arg-24, the remaining N-terminal domain secondary structure includes 5 alpha-helices, and a C-Terminal domain consisting of 4 alpha helices which is consistent with the structure found in other AcSps(Wen et al. 2018).

**Figure 4:**
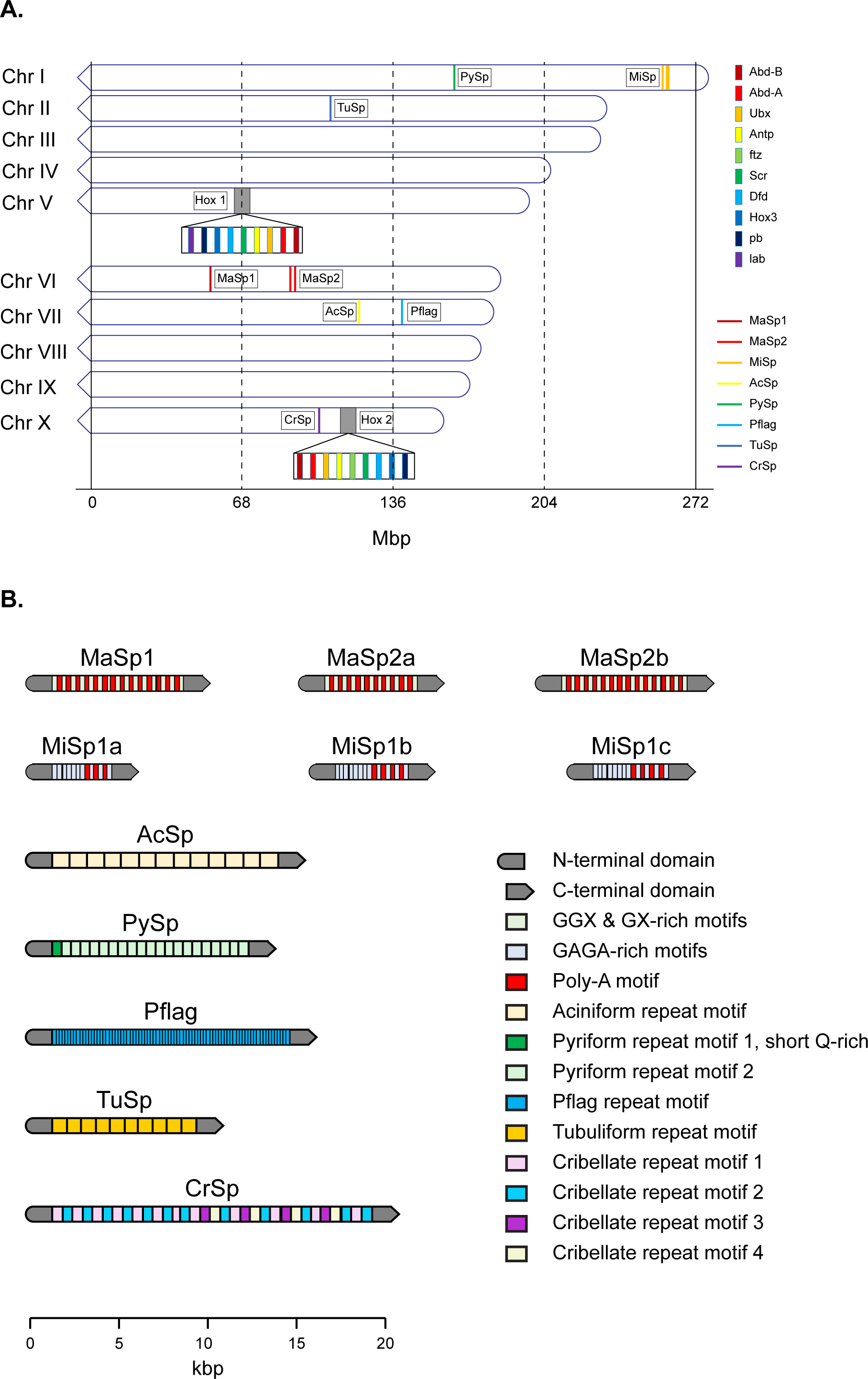
Gene Annotations. A) Gene loci for spidroins and *hox* gene clusters. B) Domain composition of identified spidoins. (Repeat region annotations are condensed for clarity.)

**Table 7.** Summary of Spidroin Gene Features.

| Spidroin | Gene Length (bp) | CDS Length (bp) | Protein Length (aa) | N-terminal Length (aa) | C-terminal Length (aa) | Signal Peptide Start | Signal Peptide Stop | Signal Peptide Probability | Signal Peptidase Cleavage Site Probability |
| --- | --- | --- | --- | --- | --- | --- | --- | --- | --- |
| Aciniform | 14,907 | 14,907 | 4,968 | 148 | 109 | Met-1 | Ser-23 | 0.9955 | 0.9056 |
| Pseudoflagelliform | 15,546 | 15,546 | 5,181 | 195 | 96 | Met-1 | Gly-29 | 0.9991 | 0.9778 |
| Cribellate | 20,198 | 20,115 | 6,704 | 874 | 296 | Met-1 | Gly-23 | 0.9997 | 0.9805 |
| Major Ampullate 2a | 7,317 | 7,317 | 2,438 | 165 | 100 | Met-1 | Gly-25 | 0.9997 | 0.9662 |
| Major Ampullate 2b | 9,141 | 9,141 | 3,046 | 195 | 109 | Met-1 | Gly-25 | 0.9687 | 0.9992 |
| Minor Ampullate 1a | 5,523 | 5,523 | 1,840 | 242 | 95 | Met-1 | Gly-23 | 0.9998 | 0.9643 |
| Minor Ampullate 2a | 6,255 | 6,255 | 2,084 | 254 | 99 | Met-1 | Gly-23 | 0.9998 | 0.9670 |
| Minor Ampullate 2b | 6,432 | 6,432 | 2,143 | 254 | 98 | Met-1 | Gly-23 | 0.9998 | 0.9670 |
| Pyriform | 13,179 | 13,179 | 4,392 | 168 | 266 | Met-1 | Gly-23 | 0.9997 | 0.9768 |
| Tubuliform | 10,146 | 10,146 | 3,381 | 179 | 346 | Met-1 | Ala-25 | 0.9993 | 0.9808 |

#### Pseudoflagelliform spidroin (Pflag)

The capture spiral of an orb-web is a composite of two types of silk(Tarakanova and Buehler 2012). For cribellate spiders such as *U. diversus*, the core fiber is the pseudoflagelliform silk made up of pseudoflagelliform spidroin (Pflag). When produced, this core fiber is coated with finely brushed cribellate silk which provides adhesive properties to the capture silk (see Cribellate spidroin).

We found a single candidate for Pflag on Chromosome 7 (**Figure 4A** and **Table 7**). We determined that the internal structure consists of 91 repeats, each of which range between 39 and 70 aa in length and are composed of two parts: a glycine-poor spacer, which is usually either 7 or 12 aa, followed by a glycine-rich repeat with variations on the motif *PSSGGXGG*. The final repeat motif in each module always terminates in a proline.

#### Cribellate spidroin (CrSp)

Cribellate silk is produced by numerous silk glands with hundreds to thousands of spigots in the cribellum. These numerous fibers are combined into a single silk which is combed into a “wooly” silk by calamistra located on the posterior legs. This wooly silk soaks into the waxy cuticle of insects by means of van der Waals interactions and hygroscopic forces and is used as the capture silk by cribellate orb-weavers(Hawthorn and Opell 2003). No full-length sequence for CrSp has been reported to date, although partial spidroin sequences have been reported for some CrSps in *Tengella perfuga*(Correa-Garhwal et al. 2018) and several *Octonoba* species(Kono et al. 2020). The whole genomic length of the *CrSp* locus on Chromosome 10 is 20,195 nt (**Figure 4A** and **Table 7**). The *U. diversus CrSp* gene was predicted to be a 2-exon gene, with a single 83 nt intron, which is consistent with what we found by manual inspection. While the entire *CrSp* locus was not spanned by single HiFi reads, we are still confident in the sequence produced, since no more than 2 reads were required to span the entire sequence. The N-terminal region of the predicted protein product consists of 874 aa and the C-terminal region consists of 296 aa (**Table 7**). The long N-terminal domain is consistent with that found in *Octonoba* spp, which were also found to have coding regions more than 2 kbp(Kono et al. 2020). We found an internal region that consists of variations on 4 repetitive motifs, with the first half of the sequence made up of motifs 1 and 2 alone, and the second half of the sequence including all 4 motifs. Motifs 1 and 3 are similar, whereas motifs 2 and 4 are distinct from each other motif (**Supplemental Figure S1)**.

#### Major ampullate spidroin (MaSp)

Draglines are produced by the major ampullate gland which produces two major ampullate spidroins (MaSp1 & MaSp2). This silk has extremely high tensile strength and elasticity and is commonly used for the primary load-bearing parts of the web such as the frame and radii. It is also the primary silk produced by spiders when they are navigating their environment(Foelix 2011). We found three candidates for major ampullate spidroin (MaSp). Based on previous work that identified multiple distinct classes of MaSps, we were able to assign one of our candidate sequences to the MaSp-1 class and the other two candidates to the MaSp-2 class. All three MaSp candidates are on Chromosome 6, although the MaSp-1 locus was located distantly from the two MaSp-2 loci (**Figure 4A**).

The MaSp-1 candidate is the single spidroin sequence we were not able to call as a complete sequence. In our annotation, the sequence appears as a two-exon gene, with the 5’ sequence and 3’ sequence found in different reading frames; however, a close inspection of the data suggests that this is not likely to be correct. We found instead that there is a large region to which the PacBio HiFi reads mapped poorly. There is consensus between the reads that indicates sequence found in the reference assembly that is not found in the reads. However, it is not clear from inspection exactly where the boundaries should be called for this region. This is likely an artifact of the assembly, since the reference was assembled from polymerase-based sequencing which is susceptible to polymerase-slippage.

The first MaSp-2 candidate, MaSp-2a, is a single exon sequence. We found that there were two distinct regions. Interestingly, the first repetitive region, which is 958 aa in length, contains mostly *GPGPQ* motifs reminiscent of the *GPGPX* motifs found in the MaSp-4 sequence recently reported in *Caerosris darwini*, but not elsewhere in the known catalog of spidroins (Garb, *et al*., 2019). The second repetitive region, which is 1,215 aa long, contains runs of poly-A and *GPX*, although *GPGPQ* repeats are also found less frequently in this region.

The second MaSp-2 candidate, MaSp-2b is also a single exon sequence. In the repetitive region, GPGPQ occurs in a few instances, but is relatively rare compared to MaSp-2a. Alternating runs of polyalanine and variations on the motif *GSGPGQQGPGQQGPGGYGPG* characterize the repetitive region. Unlike the case of the first MaSp-2 candidate, MaSp-2b does not have two distinct repetitive regions.

#### Minor ampullate spidroin (MiSp)

Minor ampullate silk has lower strength, but greater extensibility, and is composed of spidroin made by the minor ampullate gland. While it is commonly used for the construction of the auxiliary spiral in orb-weavers, it is used for prey wrapping by cob-weavers(Vienneau-Hathaway et al. 2017). We found three candidates for minor ampullate spidroin (MiSp). All three MiSp loci were located near one another on Chromosome 1 (**Figure 4A**).

The first candidate, MiSp-1 is a single exon sequence (**Table 7**). There are three repetitive regions in MiSp-1, separated by short spacers. Previous work in *Araneus ventricosus* and the cobweb weaving spiders *Latrodectus hesperus*, *L. tredecimguttatus*, *L. geometricus*, *Steatoda grossa*, and *Parasteatoda tepidariorum* has suggested that MiSp length and sequence are conserved(Chen et al. 2012; Vienneau-Hathaway et al. 2017); however, while the spacers we observed shared some sequence similarities, such as the presence of serine, threonine, and valine residues, the lengths of the spacers observed in *U. diversus* are much shorter.

The second and third candidates, MiSp-2a and MiSp-2b, shared a nearly identical amino acid composition, which was slightly different from that of MiSp-1. Both are single exon sequences (**Table** 7). Their hydrophobicity profiles are also slightly different from MiSp-1.

#### Pyriform spidroin (PySp)

Pyriform silk serves as an adhesive compound used to adhere silk lines to one-another, or to substrate that holds the web(Foelix 2011). We found one single exon candidate pyriform spidroin (PySp) on Chromosome 1 (**Figure 4A**), which is consistent in size with a prior PySp sequence reported from *Araneus ventricosus*(Wang et al. 2019a). We found that the internal repetitive region was preceded by a Q-rich N-linker region. 19 tandem repeat motifs, ranging from 188 - 196 aa were found.

#### Tubuliform spidroin (TuSp)

Tubuliform silk is used to encase the egg sac and is spun from tubuliform glands. We found a single candidate for tubuliform spidroin (TuSp) on Chromosome 2 (**Figure 4A** and **Table 7**) which is a single exon We found an internal region that was composed of 10 repeats, ranging from 262 aa - 302 aa in length. This is consistent with other reported TuSp repeats, which have been observed between 176 and 375 residues, although it seems that the typical TuSp module is repeated 15 to 20 times(Wen et al. 2017). The N-terminal region has 3 Cys residues: Cys-21, Cys-52, and Cys-132. Other TuSp N-terminal sequences have been reported with 2 Cys residues(Wen et al. 2017), however Cys-21 is expected to be removed during signal peptide cleavage. After cleavage, the N-terminal domain contains five predicted alpha helices. Cys-52 and Cys-132 are found in alpha helix 1 and alpha helix 4 after cleavage, which is also where the AcSp Cys residues are found. This conservation suggests functionality.

### Whole Genome Duplication

Gene duplication acts as a primary mode of evolutionary diversification by providing new genetic material that serves as a reservoir for subfunctionalization and neofunctionalization under selective pressure(Ohno 2013; Sémon and Wolfe 2007; Zhang 2003). In previous studies in spiders, evidence of whole genome duplication has been reported, including the presence of multiple copies of *Hox* genes(Sheffer et al. 2021; Cerca et al. 2021; Clarke et al. 2015, 2014) and the expansion of silk genes and chemosensory genes(Cerca et al. 2021). We analyzed the *U. diversus* genome to identify signatures of whole genome duplication.

Evidence for whole genome duplication can be ascribed by assessing the number of *Hox* gene clusters(Garcia-Fernàndez and Holland 1994). We used published *Hox* gene sequences from the spider *Parasteatoda tepidariorum* as query sequences for BLAST searches against the *U. diversus* genome, identifying two *Hox* gene clusters on Chromosome 5 and Chromosome 10 (**Figure 4A**), which were found to retain the expected order of *Hox* genes(Pace et al. 2016). Each cluster was missing a single *Hox* gene; however, the specific missing gene was different for each of the clusters. Cluster A on Chromosome 5 is missing the *fushi tarazu* (*ftz*) gene sequence, while Cluster B on Chromosome 10 is missing the *labial* gene sequence. Our findings are consistent with the discovery of two *Hox* gene clusters in *P. tepidariorum*(Schwager et al. 2017). In the genome of *T. antipodiana*, two *Hox* clusters were found(Fan et al. 2021), including one complete cluster on chromosome 12 which included a copy of all 10 expected *Hox* genes, and a second cluster on chromosome 8 which was found to be missing *abdominal-A*, *Hox3*, *ftz*, and *Ultrabithorax*.The presence of multiple *Hox* clusters in the *U. diversus* genome adds further support to an ancient, ancestral whole genome duplication.

We searched for evidence of synteny between pseudochromosomes using AnchorWave(Song et al. 2022). The initial identification of syntenic blocks was quite ubiquitous across all 10 pseudochromosomes, but with large gaps between mRNAs and/or with very large interanchor distances. To constrain these results, we chose to include only mRNAs where the number of missing mRNAs between anchors was 4 or less. We made this allowance to account for the fact that we expect there to be significant loss of copies of duplicated genes(Ohno 2013; Sémon and Wolfe 2007; Hakes et al. 2007). This resulted in retaining 196 blocks of at least 2 mRNAs (**Figure 5**). Considering the conservative nature of our analysis, this line of evidence provides further support for an ancient duplication event. However, considerable reorganization has occurred since the duplication event, an observation also made in mammalian genomes, and potentially associated with their successful adaptation to diverse environments(Waters et al. 2021; White 1978).

**Figure 5:**
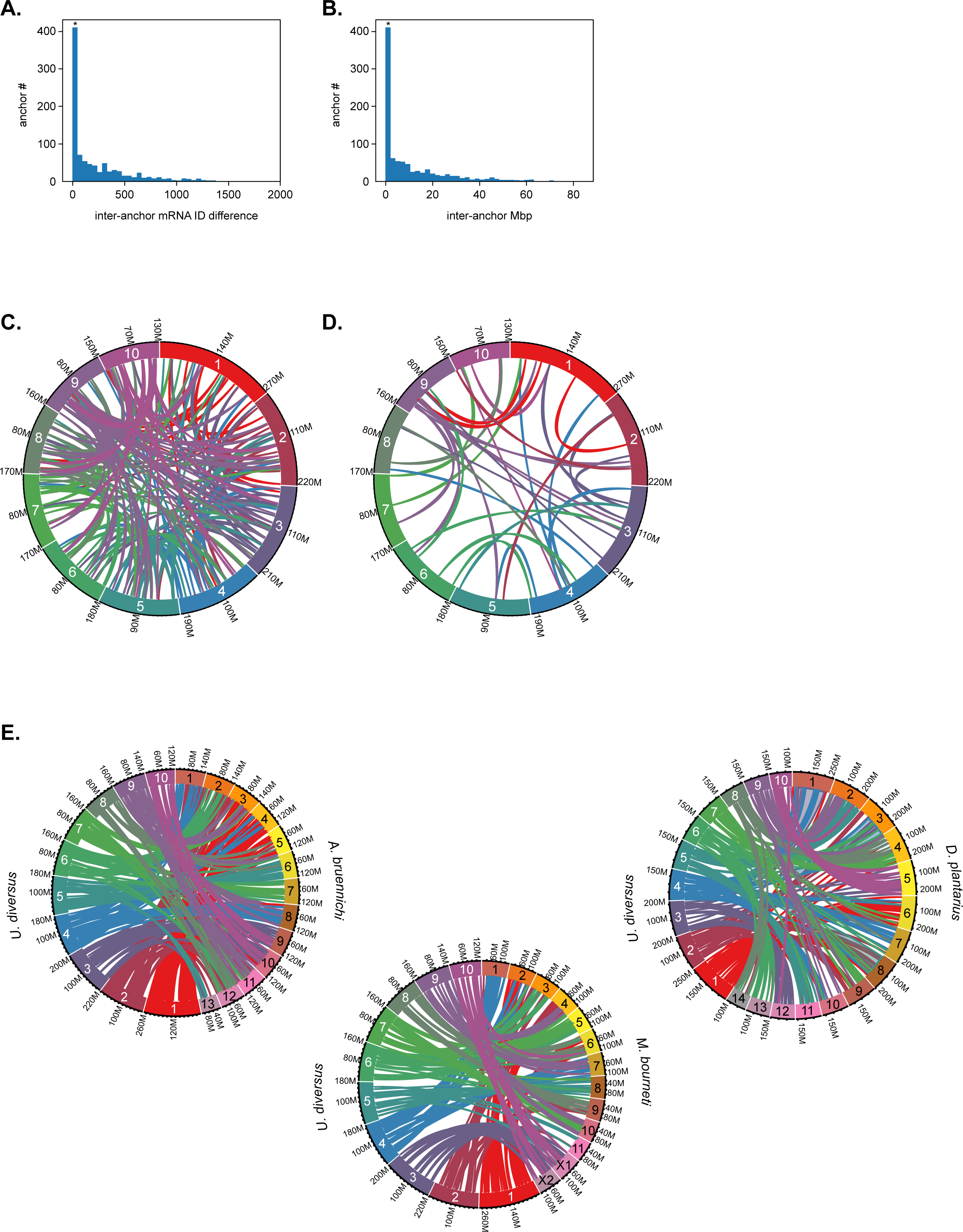
Synteny and Chromosomal Rearrangements. A) Inter-anchor mRNA ID difference distribution of syntenic blocks identified by AnchorWave analysis. Each syntenic block is defined by ORF or inter-ORF anchors. All ORFS are numerically annotated in consecutive order from scaffold 1 through scaffold 10. Inter-anchor mRNA ID difference is defined as the difference in these numerical ORF IDs between consecutive ORF anchors. If the distance equals 1, it means the two anchors are consecutive ORFs within the block. Asterisk indicates syntenic blocks used in **E**. B) Inter-anchor Mbp difference distribution of syntenic blocks identified with AnchorWave analysis. Inter-anchor difference was calculated as the base-pair distance between consecutive ORF anchors within a syntenic block. Asterisk indicates syntenic blocks used in **E**. C) Ribbon plot of all AnchorWave-defined syntenic blocks shared between chromosomal scaffolds. D) Ribbon plot of filtered AnchorWave-defined syntenic blocks shared between chromsomal scaffolds. Only blocks consisting of consecutive ORF anchors < 4 mRNA IDs apart are plotted. E) Ribbon plot of filtered AnchorWave-defined syntenic blocks shared between *U. diversus* and *A. bruennichi, M. bourneti,* and *D. plantarius* chromsomal scaffolds. Only blocks consisting of consecutive ORF anchors < 4 mRNA IDs apart are plotted.

We also compared the synteny of our *U. diversus* pseudochromosomes with those from *Argiope bruennichi*, an Araneid orb-weaver, *Meta bourneti* a Tetragnathid orb-weaver, as well as *Dolomedes plantarius* a Pisaurid (**Figure 1**). While the syntenic blocks of some of the chromosomes seem split between multiple chromosomes across the species, certain chromosomes, or pairs of chromosomes, have nearly exclusive synteny between species. *A. bruennichi* chromosomes 6, 7, and 8 share nearly exclusive syntenic blocks with *U. diversus* chromosomes 5, 6, and 4, respectively. This shared synteny with *U. diversus* chromosomes 5 and 6 is also observed for chromosomes 8 and 5 from *M. bourneti*, however U. diversus chromosome 4 is split between M. bourneti chromosomes 1 and 7. *U. diversus* chromosome 1 shares considerable synteny with *A. bruennichi* chromosomes 3-5, while *U. diversus* 2 is biased for shared synteny with A. bruennichi chromosomes 1-2 and 11-12. Even though *D. plantarius* diverged more recently from *U. diversus* (**Figure 1**), there appears to a greater degree of chromosomal rearrangement between these two species. However, two outliers are *U. diversus* 3 and 10 which share nearly exclusive synteny with *D. plantarius* 12 and 5, respectively. The large degree of synteny between *U. diversus* 3 and 10 with *M. bourneti* X1 and X2 is a strong indication that these two chromosomes are the sex chromosomes for *U. diversus*.

### Sex Chromosomes

The most common and likely ancestral system of chromosomal sex determination in spiders is ♂X_1_X_2_/♀X_1_X_1_X_2_X_2_.(Sember et al. 2020) However, spiders exhibit a diversity of sex determining systems, with some including Y chromosomes, and others possessing up to 13 X chromosomes(Král et al. 2019). Usually, these sex determining systems are determined by karyotyping(Sember et al. 2020; Král et al. 2019) (**Figure 3**). The genetic basis of sex determination and chromosomal dosage compensation are unknown for spiders. Determining the genetic identity of X chromosomes has been challenging, in part due to significant levels of shared synteny between sex chromosomes and autosomes(Sember et al. 2020) (**Figure 5**), as well as a paucity of spider genomes with chromosome-level scaffolds. X chromosome scaffolds from more fragmented genomes have been identified by quantifying the relative difference in read depth from sperm with or without the X chromosomes(Bechsgaard et al. 2019). Recently, the X chromosomes for *A. bruennichi* (also ♂X_1_X_2_/♀X_1_X_1_X_2_X_2_) were identified through disparities in read coverage of X chromosomes between males and females(Sember et al. 2020). In principle, because males have only one copy of each X chromosome, the average read depth for scaffolds from these chromosomes should be half that of autosomes. To determine the sex chromosome in *U. diversus*, we assembled an Illumina short-read library from a single male spider and mapped the reads onto the 10 assembled pseudochromosomes (**Figure 6**).

**Figure 6:**
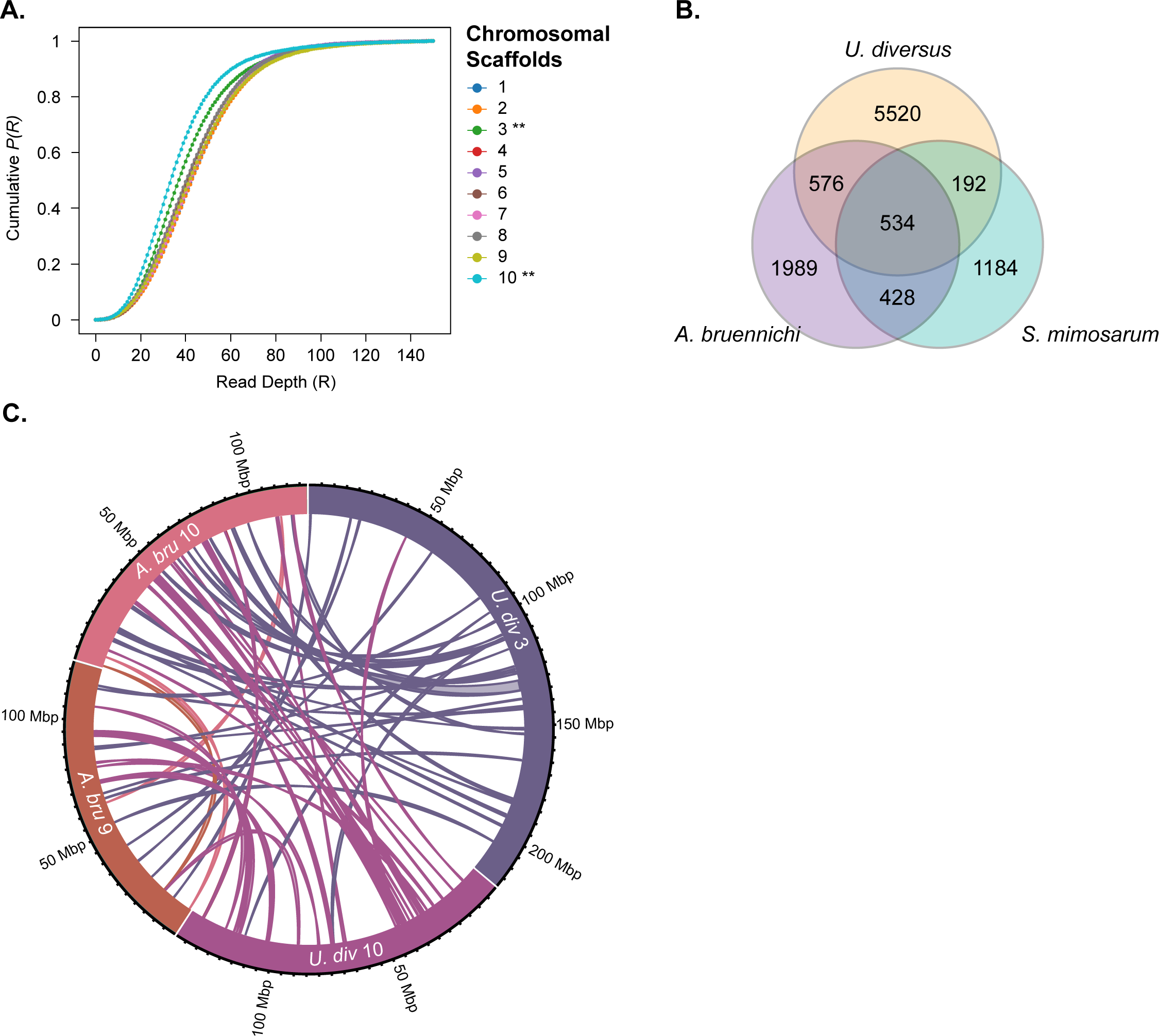
Sex Chromosomes. A) Read depth of Illumina reads from a male spider aligned to the chromosomal scaffolds. Scaffolds 3 and 10 (asterisks) exhibited lower read depth than other scaffolds. B) Venn diagram of shared sex-associated genes identified in *U. diversus*, *S. mimosarum*, and *A. bruennichi*. C) Ribbon plot of shared synteny between predicted X chromosomes from *U. diversus* and *A. bruennichi*.

While 8 of the 10 pseudochromsomes had a median read depth of 40 ± 2, pseudochromosomes 3 and 10 were outliers, with read depths of 36 and 33, respectively. If these pseudochromosomes were exclusively unique X chromosomes, the expected read depth would have been ∼20. However, as observed in other species(Sember et al. 2020) and our own (**Figure 5**), orthologous autosomal regions should decrease the expected depth disparity. The higher than expected read depth could also be due to mis-assembly of these pseudochromosomes, however very little linkage was observed between pseudochromosomes 3 and 10 in the Hi-C data (**Figure 3**). Despite these caveats, the lower median read depth in males for pseudochromosomes 3 and 10 is a strong indicator these likely represent the two X chromosomes for *U. diversus*.

Prior work with *Stegodyphus mimosarum* (also ♂X_1_X_2_/♀X_1_X_1_X_2_X_2_) identified sex-linked scaffolds based on lower read depth of sperm lacking X chromosomes(Bechsgaard et al. 2019). When genes identified on these X-linked *S. mimosarum* scaffolds were mapped on to the *U. diversus* pseudochromosomes (**Table 5**), 62% of these genes mapped onto pseudochromosomes 3 and 10 (**Figure 6B**). This large fraction of predicted X-linked genes between two distantly related species of spiders is a strong indicator that not only are pseudochromosomes 3 and 10 likely to be the X chromosomes, but that the genetic composition of these chromosomes has remained fairly stable amongst spiders. Since two X chromosomes were recently identified in *A. bruennichi*, we compared the genetic composition (**Supplemental Table S6**) and synteny between the X chromosomes identified in both species (**Figure 6**). In addition to shared X-linked genes (**Supplemental Table S6**, **Figure 6B**), *A. bruennichi* scaffolds 9 and 10 appear to share considerable synteny with *U. diversus* pseudochromosomes 3 and 10 (**Figure 6C**), while sharing little synteny with the autosomes of the other species (**Figure 5E**). This also appears to be true of the sex chromosomes X1 and X2 from *M. bourneti*, however this genome is not currently annotated. When chromosomal rearrangements have occurred, they appear to have been confined to rearrangements between sex-chromosomes. The sex chromosomes themselves share little synteny with each other (**Figure 5**), which indicates they are not the result of an ancient duplication, but there appears to be selective pressure to ensure that when chromosomal rearrangements do occur, that they occur between the sex chromosomes. However, some syntenic blocks are shared with autosomes. One of these syntenic blocks is the *Hox* gene cluster located on chromosome 10 for both *U. diversus* and *A. bruennichi*. The presence of a *hox* cluster on a sex chromosome was surprising since these genes play critical roles in development. Therefore, either dosage compensation is needed in males, or dosage disparity between males and females plays a role in developmental sexual dimorphism.

In insects, the primary sex chromosome dosage sensor is *sex lethal* (*sxl*), which then triggers a cascade of sex-defining signaling events leading to sexually dimorphic expression of genes and/or splice variants. While no *sxl* homologue has been found in spider genomes (including *U*. *diversus*), other genes involved in sexual dimorphism, such as *doublesex* (*dsx*) are present. Thus, the mechanism spiders use for sensing X:autosome ratio differences remains unknown, but relevant genes are likely shared between *U. diversus*, *A. bruennichi*, and *S. mimosarum*. Of the 534 shared sex-linked genes in these three species, 14 are predicted to be DNA/RNA-binding, and may play a role in sex-determination. The X-linked genes shared between these three species (**Supplemental Table S6**) will be a resource for comparative analysis to identify conserved genes that serve as sex-specifying triggers for spiders. Uncovering how spiders perform sex-linked dosage compensation can not only illuminate how arthropods evolved different sex-determining systems, but also how dosage compensation has evolved independently in numerous animals.

## Conclusions

Here, we present a high-quality chromosome-level genome and complementary transcriptome assembly of the hackled orb-weaver *Uloborus diversus*. The 2.15 Gbp draft genome assembly comprises 1,586 scaffolds, including 10 pseudochromosomes that contain 1.9 Gbp (88%) of the total assembly, comparable to the estimated genome size (1.98 Gbp) predicted by GenomeScope2 and contains the vast majority of highly conserved orthologs (94.1% complete, with 88.6% complete and in single copy) as estimated by BUSCO. We predicted a total of 44,408 protein-coding gene models with a BUSCO completeness of 86.7%. Despite the aforementioned technical hurdles, the contiguity and completeness of this assembly, along with the recovery of a complete catalog of full-length spidroin gene sequences, demonstrates the utility of using multiple complementary sequencing technologies for large, repetitive, and highly heterozygous genomes.

The repetitive nature and length of spidroin genes have posed a technical challenge for identifying and reporting full-length sequences. However, it is exactly these qualities that lend spidroins their unique mechanical properties(Malay et al. 2017; Rising et al. 2005; Li et al. 2017); underscoring the need for accurate assemblies. Recent studies leveraging single molecule, long read sequencing technology have predicted longer spidroin sequences than those using PCR approaches(Kono et al. 2019). Here, we used ONT and PacBio HiFi reads to achieve a complete catalog of full-length spidroin sequences for *Uloborus diversus*. The ability to recover full-length sequences for this family of genes is an indication of the high quality of the assembly.

All current models of chromosomal dosage compensation are based on single-sex chromosome animals, however multiple sex-chromosome systems exist in both vertebrates and invertebrates(Yoshido et al. 2020; Rens et al. 2007). Spiders exhibit considerable morphological and behavioral sexual dimorphism that is based on a multiple-sex chromosome system. Understanding the genetic underpinnings of spider sexual development will contribute to a fuller understanding of how chromosomal sex determination can evolve independently in different species. Here we provide evidence for the identities of sex chromosomes in *U. diversus* and leverage this information to identify 14 candidate DNA-binding genes that are shared between three divergent species of spiders.

Our genome will facilitate comparative studies and meets a specific need in the field for a greater representation of genomes from the UDOH+RTA clade that represent nearly half of all known spider species (**Figure 1**)(Garb et al. 2018). We expect that the highly contiguous draft genome and transcriptome datasets we produced for *U. diversus* will serve as a valuable resource for continuing research into the evolution, development, and physiology of spiders, as well as a vital tool to study the genetic basis of orb-weaving behavior. While a handful of spider genomes have been published, all orb-weaving genomes have been from ecribellate Araneid spiders, with no representative genomes from the cribellate families Uloboridae, Deinopidae, Oecobiidae, or Hersiliidae. Improved knowledge of genomes from these families, combined with behavioral and cellular analyses of orb-weaving behavior, will offer a crucial foundation for understanding how and when orb-weaving evolved.

## Materials and Methods

### Sample Collection and Husbandry

We collected spiders of the species *Uloborus diversus* from the ancestral lands of the Ramaytush, in Half Moon Bay, California, USA. We collected colony founders from a single greenhouse during several trips between 2016 and 2019 and transported them to custom-fabricated habitats in an on-campus greenhouse at Johns Hopkins University. We later transferred experimental animals from the on-site greenhouse to custom-fabricated habitats in the laboratory until required for experiments. We fed all animals alternately *Drosophila melanogaster* or *Drosophila virilis* once per week.

### Karyotyping

We soaked embryos soaked in Grace’s insect medium (Gibco) containing 0.1% colchicine for 2 hours. We then added an equal volume of hypotonic solution. After 15 min, we transferred the embryos to a 3:1 ethanol:acetic acid solution for 1 hour. After fixing, we transferred embryos to gelatin-coated microscope slides and dissociated them in a drop of 45% acetic acid. We used siliconized coverslips to squash the dissociated tissue and briefly froze them in liquid nitrogen. After removing the slides from LN2, we immediately removed the coverslips with a razor blade and transferred the slides as quickly as possible to 95% ethanol. We then performed a step-down series from 95% ethanol to 70%, 35%, and finally to Grace’s insect medium to return the tissue to an aqueous solution. We then transferred the slides to a 1ug/mL DAPI solution. After a 10-minute incubation, we transferred the slides to de-ionized water to rinse and mounted coverslips with a drop of Vectamount (Vector Laboratories, Burlingame, CA, USA).

### RNA Extraction, Library Preparation, and Sequencing

We extracted total RNA from multiple samples: a whole adult female, a whole adult male, adult female prosoma and opisthosoma, adult male prosoma and opisthosoma, pooled legs from both the adult female and adult male dissections, a 4th instar female, and approximately 30 pooled 2nd instars. We used the Qiagen RNeasy Mini Kit (Qiagen, Hilden, Germany) to extract total RNA, following the manufacturer’s protocol. We estimated the quality and quantity of total RNA using a NanoDrop One Microvolume UV-Vis Spectrophotometer (ThermoFisher Scientific, Waltham, MA, USA). Before library preparation, we also measured the quality, quantity, and fragment length of our total RNA using a TapeStation 4200 System with RNA ScreenTape and reagents (Agilent, Santa Clara, CA, USA). We prepared barcoded, directional, paired-end RNA-seq libraries with the NEBNext Ultra II Directional RNA Library Prep Kit for Illumina using the NEBNext Poly(A) mRNA Magnetic Isolation Module. We submitted the resulting libraries to the Johns Hopkins Genomics Core Resources Facility to be sequenced on an Illumina HiSeq 2500 Sequencing System with 150 bp paired-end chemistry.

### Genomic DNA Extraction, Library Preparation, and Sequencing

Prior to extraction of DNA, we withdrew food for 3 days to minimize the potential contribution of contaminating DNA from dietary sources. We extracted high molecular weight (HMW) DNA using the QIAgen MagAttract HMW DNA kit. Prior to HMW purification, we followed the manufacturer’s protocol for disruption/lysis of tissue. We avoided fast pipetting and prolonged vortexing to minimize shearing of DNA. We flash froze adult spiders in liquid N_2_ and crushed them with a pellet pestle (Fisher, 12-141-364) in a Protein LoBind tube (Eppendorf, 022431081) containing 220 uL of Buffer ATL. We then added 20 uL Proteinase K and briefly vortexed the sample. We next incubated the sample overnight at 56C with 900 rpm shaking on a ThermoMixer C (Eppendorf, 5382000023). After the overnight incubation, we then briefly centrifuged the sample to spin down condensate on the tube. We next transferred 200 uL of lysate to a fresh 2 mL sample tube and followed the manufacturer’s protocol for manual purification of HMW DNA from fresh or frozen tissue. We estimated DNA quality using a NanoDrop One Microvolume UV-Vis Spectrophotometer and quantified DNA using a Qubit 4 Fluorometer (ThermoFisher) with a Quant-iT dsDNA HS Assay Kit. We also measured DNA quality, quantity, and fragment length distributions using the Agilent TapeStation 4200 System with Genomic DNA ScreenTape and reagents before proceeding to library prepration. A typical preparation from a 20 mg spider yielded 8.5ug of DNA.

#### Illumina Sequencing

For Illumina sequencing, we extracted genomic DNA from a single, whole, unmated penultimate stage female to minimize the potential contribution of extraneous haplotypes from stored sperm after mating events. We submitted the HMW gDNA to the Johns Hopkins GCRF, where they prepared a PCR-free library of approximately 400 bp DNA insert size using the Illumina TruSeq PCR-Free High Throughput Library Prep Kit (San Diego, CA, USA), according to the manufacturer’s protocol. They then sequenced the prepared library on an Illumina NovaSeq 6000 Sequencing System with 150bp paired-end chemistry.

#### ONT Sequencing

For ONT sequencing, we extracted HMW genomic DNA from 3 adult females. We prepared sequencing libraries using the Ligation Sequencing Kit (SQK-LSK109) (Oxford Nanopore Technologies, UK), according to the manufacturer’s protocols. Third party reagents we used during library preparation included: New England Biolabs (New England Biolabs, Ipswitch, MA, USA) NEBNext End Repair/dA-Tailing Module (E7546), NEBNext FFPE DNA Repair Mix (M6630), and NEB Quick Ligation Module (E6056). We then sequenced the libraries, using ONT

R.9.4.1 flowcells (FLO-PRO002) on an ONT PromethION sequencing platform. We then used ONT’s Albacore basecalling software v.2.0.1 (RRID:SCR_015897) to basecall the raw fast5 data.

#### PacBio HiFi Sequencing

For PacBio sequencing, HMW DNA was extracted from a single adult female spider provided to Circulomics (Baltimore, MD, USA). They extracted DNA using a modified protocol with the Nanobind Tissue Kit (Circulomics, #NB-900-701-01). Briefly, they froze and crushed a single, adult female spider with a pellet pestle (Fisher, #12-141-364) in a Protein LoBind tube (Eppendorf, #022431081) containing 200 uL of Buffer CT. The crushed spider was centrifuged at 16,000 x g at 4 C for 2 min. The supernatant was discarded, and the pellet was resuspended in 500 ul Buffer CT and the mixture was transferred to a 2.0 mL Protein LoBind tube (Eppendorf # 022431102). The suspension was spun again at 16,000 x g at 4 C for 2 min and the supernatant discarded. The spider tissue pellet was combined with 20 ul Proteinase K and 150 ul Buffer PL1 and resuspended by pipetting with a P200 wide bore pipette tip. The tissue was incubated on a ThermoMixer at 55 C with 900 rpm mixing for 1 hour. After lysis, 20 ul RNaseA was added, and the lysate was mixed by pipetting with a P200 wide bore pipette tip. The lysate was incubated at RT for 3 min. After RNaseA incubation, 25 ul Buffer SB was added, the lysate was vortexed 5 x 1 sec pulses, and then centrifuged at 16,000 x g at 4 C for 5 min. The supernatant (∼200 ul) was transferred to a 70uM filter (Fisher # NC1444112) set in a new 1.5 mL Protein LoBind tube (Eppendorf # 022431081). The tube with the 70 uM filter was spun on a mini-centrifuge (Ohaus # FC5306) for 1 sec and then the filter was dicarded. 50 ul Buffer BL3 was added to the cleared lysate and the tube was inversion mixed 10X. The tube was then incubated on a ThermoMixer at 55 C with 900 rpm mixing for 5 min. After incubation, the tube was allowed to come to RT, which took about 2 min. The tube was spun for 1 sec on a mini-centrifuge to spin down condensate from the lid. One 5 mm Nanobind disk was added to the tube followed by 250 uL isopropanol and then the tube was inversion mixed 5X. The tube was then rocked on a platform rocker (ThermoScientific # M48725Q) at RT and max speed for 30 min. The DNA-bound Nanobind disk was washed according to handbook directions with one 500 ul CW1 wash and one 500 ul CW2 wash. The tube with the disk was tap spun for 2 x 1 sec to dry the disk. The DNA was eluted with 50 ul Buffer EB and incubated at RT overnight. The next day, the eluate was pipette mixed with a standard bore pipette tip 5x and then quantitated with Nanodrop and Qubit dsDNA BR assay and then sized by pulsed-field gel electrophoresis.

We then submitted the DNA sample to the University of Maryland School of Medicine Genomics Core Facility for PacBio HiFi sequencing. There, they size selected the DNA using a Safe Science BluePippin with a 9kb high-pass cutoff. They prepared the sequencing library using the Express v2 kit, according to the standard protocol for preparing HiFi sequencing libraries. They then sequenced the library on a PacBio Sequel II 8M SMRT Cell using a 30 hour HiFi run mode and processed using SMRT Link v.9.0 software.

#### Dovetail Chicago and Dovetail Hi-C Sequencing

To further improve the *U. diversus* genome assembly, we used proximity ligation-based sequencing techniques to scaffold intermediate versions of our assembly. We provided 19 spider specimens to Dovetail Genomics (Scotts Valley, CA, USA) for Chicago and Hi-C library preparation as previously described(Putnam et al. 2016). They prepared a Chicago library using 15 pooled adult females and a Hi-C library using 4 pooled adult females. They sequenced both the prepared Chicago and Dovetail Hi-C libraries on an Illumina HiSeq X sequencing platform on 1 flowcell.

### DNA-seq and RNA-seq QA/QC

For Illumina, we examined read quality using FastQC(Andrews 2010) v.0.11.9 (RRID:SCR_014583). For DNA-seq data, we determined that, due to high quality of reads and absence of adapter sequences, no further processing would be required and proceeded to assembly with raw read data.For RNA-seq data, we used TrimGalore(Krueger et al. 2021) v.0.4.2 (RRID:SCR_011847) to apply quality filtering and remove adapter sequences from the FASTQ files. We performed additional filtering for quality with Trimmomatic(Bolger et al. 2014) v.0.33 (RRID:SCR_011848) . For ONT, reads shorter than 3 kbp were discarded. The length-filtered ONT long reads were used in downstream assembly.

### Genome Size, Heterozygosity, and Unique Sequence Estimation

Prior to assembly, we used Jellyfish(Marçais and Kingsford 2011) v.2.2.4 (RRID:SCR_005491) to count the frequency of canonical 21-mers in our Illumina sequencing data. We used the resulting sorted *k*-mer frequencies vs counts histogram as input to GenomeScope(Ranallo-Benavidez et al. 2020; Vurture et al. 2017) v.2.0 to estimate genome size, heterozygosity, and repetitiveness.

### Recovery of Mitogenome

We used Novoplasty(Dierckxsens et al. 2016) v.4.2 (RRID:SCR_017335) to generate a complete circularized mitochondrial sequence using raw Illumina read data. The mitochondrial sequences of several spider species were used to provide seed sequences (**Supplemental Table S3**). The resulting mitogenome sequences assembled by Novoplasty were compared for consensus. The consensus mitogenome was uploaded to the MITOS 2 web server(Bernt et al. 2013) for annotation. The CGView web server(Stothard and Wishart 2005) (RRID:SCR_011779) was used to visualize the annotated mitogenome.

### Nuclear Genome Assembly

#### De novo Nuclear Genome Assembly with MaSuRCA

Illumina reads were assembled into contigs and the resulting contigs were scaffolded with ONT long reads using the MaSuRCA assembly pipeline(Zimin et al. 2013, 2017) v.3.4.2 (RRI:010691). We used default settings, including the default CABOG contigging module in lieu of the Flye assembler. The resulting genome assembly is referred to as *U. diversus* v.1.0.

To improve the assembly, we used *Rascaf*(Song et al. 2016) v.2016-11-29 to scaffold with Illumina RNA-seq read data. The resulting genome assembly is referred to as *U. diversus* v.1.1. To reduce redundancy in the assembly due to the presence of alternative haplotigs, we used Pseudohaploid with default settings. The resulting genome assembly is referred to as *U. diversus* v.1.2

#### De novo Nuclear Genome Assembly with PB-IPA

We used PacBio’s Improved Phased Assembly (IPA) HiFi Genome Assembler with default settings, specifying a genome size of 1.9 Gbp, to assemble the HiFi reads. The resulting genome assembly is referred to as *U. diversus* v.2.0.

#### Merging MaSuRCA and PB-IPA Assemblies with SAMBA

We used the SAMBA tool distributed with MaSuRCA to merge the MaSuRCA assembly, *U. diversus* v.1.2, and the PB-IPA assembly, *U. diversus* v.2.0. The resulting genome assembly is referred to as *U. diversus* v.3.0.

### Scaffolding Assemblies with *HiRise*

The initial *U. diversus* v.1.2 draft assembly obtained using a combination of MaSuRCA, Rascaf, and Pseudohaploid was provided to Dovetail Genomics in FASTA format. The resulting genome assembly is referred to as *U. diversus* v.1.3.

The merged MaSuRCA and PB-IPA assembly, *U. diversus* v.3.0, was provided to Dovetail Genomics in FASTA format. The resulting genome assembly is referred to as *U. diversus* v.3.1.

### Genome Assembly Metrics and Assessments

For each assembly, completeness was estimated with Benchmarking Universal Single-Copy Orthologs (BUSCO)(Seppey et al. 2019; Simão et al. 2015; Waterhouse et al. 2018) v.5.2.1 (RRID:SCR_015008) using the arachnida_odb10 database(Kriventseva et al. 2019). Contiguity of each assembly was evaluated for comparison using Quast(Gurevich et al. 2013) v.5.0.2 (RRID:SCR_001228).

### Genome-Guided Transcriptome Assembly

Cleaned and trimmed Illumina RNA-seq reads were aligned to the genome using *HISAT2*(Kim et al. 2019) v.2.2.1. We then used the Trinity assembler^43^ v.2.12.0 to produce a genome-guided transcriptome assembly [--CPU 60 –max_memory 200G –genome_guided_max_intron 20000 – SS_lib_type RF –include_supertranscripts –verbose]. We used TransDecoder(Haas et al. 2013) v.5.5.0 with default settings, including homology searches using both BlastP(Altschul et al. 1990; Altschul 1997) against a SwissProt UniProt database(The UniProt Consortium 2019) as well as the Pfam database(Mistry et al. 2021) v.32, as ORF retention criteria.

### Repeat Annotations

To characterize the repeat elements in the *U. diversus* genome, we generated a custom *de novo* repeat library using RepeatModeler(Flynn et al. 2020) v.2.0.2 with default parameters. We used RepeatMasker(Tarailo□Graovac and Chen 2009) v.4.1.2 to screen and mask repeat and low-complexity regions of the genome with the Dfam consensus(Storer et al. 2021) v.3.4 and RepBase RepeatMasker Edition(Bao et al. 2015) v.2018-10-26 repeat libraries.

### Annotation of Protein Coding Genes

We performed gene annotation using the BRAKER 2 pipeline(Hoff et al. 2016; Brůna et al. 2021; Lomsadze 2005; Lomsadze et al. 2014; Stanke et al. 2006, 2008; Gotoh 2008; Li et al. 2009; Barnett et al. 2011; Iwata and Gotoh 2012; Buchfink et al. 2015; Hoff et al. 2019; Brůna et al. 2020) v.2.1.6 with RNA-seq evidence and protein homology evidence based on a custom library of spider sequences obtained from NCBI. BRAKER2 uses RNA-seq data to produce intron hints for training the *ab initio* gene prediction program AUGUSTUS(Stanke et al. 2006, 2008) on a species-specific model. This species-specific model is then used in conjunction with RNA-seq data to predict protein coding genes. The bam file previously generated in transcriptome assembly and analysis was passed to BRAKER2, which was run with default settings.

### Annotation of Non-Coding RNAs

We used tRNAscan-SE(Chan and Lowe 2019; Chan et al. 2021) v.2.0.7 with default settings to predict tRNAs. We then used Barrnap(Seeman 2018) v.0.9 (RRID:SCR_015995) with default settings to predict rRNAs.

### Functional Annotation

We started the annotation of predicted genes used the BLAST+ blastp algorithm. First, we obtained the longest coding sequence for each gene predicted by BRAKER2. We then used the EMBOSS(Rice et al. 2000) v.6.6.0.0 Transeq tool to translate and trim the coding sequences. Once translated and trimmed, we used the BLAST+ v.2.10.1+ Blastp tool to search against the UniProt SwissProt database with an e-value cutoff of 1e-10. We used InterProScan(Jones et al. 2014; Quevillon et al. 2005) (RRID:SCR_005829) to predict motifs, domains, and gene ontology (GO)(Ashburner et al. 2000; The Gene Ontology Consortium et al. 2021) terms (RRID:SCR_002811), as well as MetaCyc(Caspi et al. 2016, 2018) and Reactome(Gillespie et al. 2022; Jassal et al. 2019) pathways,using the following analyses: CDD(Lu et al. 2020) v.3.18, Coils v.2.2.1, Gene3D(Lewis et al. 2018) v.4.3.0, Hamap(Pedruzzi et al. 2015) v.2020-05, MobiDBLite(Necci et al. 2017) v.2.0, PANTHER(Mi et al. 2019) v.15.0, Pfam(Mistry et al. 2021) v.34.0, PIRSF(Wu 2004) v.3.10, PIRSR(Chen et al. 2019a) v.2021-02, PRINTS(Attwood 2003) v.42.0, ProSitePatterns(Sigrist 2002; Sigrist et al. 2012) v.2021-01, ProSiteProfiles(Sigrist 2002; Sigrist et al. 2012) v.2021-01, SFLD(Akiva et al. 2014) v.4, SMART(Letunic and Bork 2018; Letunic et al. 2021) v.7.1, SUPERFAMILY(Pandurangan et al. 2019; Gough et al. 2001) v.1.75, and TIGRFAM(Haft et al. 2012; Selengut et al. 2007; Haft 2003, 2001) v.15.0.

### Spidroins

#### Identification of Spidroin Candidate Sequences

We identified *Uloborus diversus* by conducting BLAST(Altschul et al. 1990; Altschul 1997) searches using the list of spidroin sequences included in **Supplemental Table S5** as queries against the assembled genome, transcriptome, and gene models predicted by BRAKER2. We looked for matches to both N- and C-terminal sequences from members of each type of spidroin, as well as to available repetitive motifs. After cross-referencing genomic coordinates with gene models and transcripts, we used JBrowse(Buels et al. 2016) to visualize mapping of Illumina RNAseq data and PacBio HiFi reads to the assembled genome. RNAseq reads were mapped to the genome with HISAT2(Kim et al. 2019), while minimap2(Li 2018, 2021) was used to map PacBio HiFi reads. Samtools(Li et al. 2009) was used to convert the resulting SAM files to BAM files, as well as to sort and index the BAM files. For each spidroin candidate, the entire sequence from start codon to stop codon, ignoring any predicted splicing, with an additional 5 kb of sequence on both the 5’ and 3’ end, was translated in all six frames using the ExPASy Translate Tool via the ExPASy web server(Gasteiger 2003) and inspected for ORFs as well as the presence of repetitive motifs characteristic of spidroins. Predicted splice sites were compared with RNAseq data. Unsupported splice sites, either by lack of evidence in the mapping of RNAseq reads or by the obvious presence of spidroin repeat motifs within the predicted intronic region, were removed from the annotations. Spidroins sequences were called based upon the preponderance of available evidence, which in some cases conflicted with the structure predicted by BRAKER2.

#### Spidroin Sequence Analysis

We used the ExPASy web server tool ProtScale to find the amino acid composition of each sequence, as well as to estimate the hydrophobicity using the the Kyte-Doolittle method(Kyte and Doolittle 1982; Gasteiger 2003). We used the PSIPRED v.4.0 tool in the UCL Bioinformatics Group’s PSIPRED Protein Analysis Workbench(Buchan et al. 2013) to predict the secondary structure of each sequence. The sequences were often too long and necessitated judicious segmentation into reasonable sequences that were short enough for analysis. In such cases, we selected natural breaks in the sequence structure, such as separating the N-terminal region from the repetitive regions, etc. We used SignalP(Teufel et al. 2022) v.6.0 to predict the presence signal peptides and signal peptidase cleavage sites in the N-terminal regions.

## Data Availability

The raw sequencing data and assembled genome presented in this study have been submitted to the NCBI BioProject database (https://www.ncbi.nlm.nih.gov/bioproject/) under accession number PRNA846873.

## Author Contributions

J.M., A.Z. and A.G. designed the research study. J.M. performed DNA purification and sample preparation for Illumina and Oxford Nanopore sequencing. J.M. performed all computational analyses, except for HiRise scaffolding (performed by Dovetail), MaSuRCA and SAMBA. A.Z. performed MaSuRCA assembly and merging with SAMBA. J.M. and A.G. analyzed the data and wrote the paper.

## Acknowledgements

We thank Circulomics Inc., particularly Kelvin Liu and Michelle Kim, for assistance in DNA extraction for PacBio HiFi sequencing. We thank the Johns Hopkins University Genomics Core, and David Mohr in particular, for Illumina sequencing and consultation. We additionally thank the University of Maryland Genomic Resource Center, and Luke Tallon specifically, for PacBio HiFi sequencing. We thank Dovetail Genomics, particularly Mark Daly and Tom Swale, for Chicago and Dovetail Hi-C library preparation and sequencing, as well as HiRise assembly scaffolding and consultation. We thank the members of the Timp Lab, in particular Winston Timp, Norah Sadowski, and Rachael Workman, for training and graciously permitting the use of their ONT PromethION and TapeStation. We thank Gordus lab members, James Taylor, Michael Schatz, Bob Johnston, Rajiv McCoy, Prashant Sharma, and Ben Matthews for helpful discussions and comments on the manuscript.

---

Table S1 - Comparison of Genome Statistics for Published Genomes.

Table S2 - Summary of Spider Genome Repeat Content.

Table S3 - Library of Annotated Genes from Spider Genomes.

Table S4 - Mitogenome Sequences Used for NOVOplasty Seeds.

Table S5 - Spidroin Protein Sequences Used in BLAST Searches.

Table S6 - Summary of Common Sex-Linked Annotations.

