## Supplementary material for "Chromosome-level genome and the identification of sex chromosomes in Uloborus diversus": Table S2

**Table S2. Summary of Spider Genome Repeat Content**

| Species | SINEs (%) | LINEs (%) | LTR elements (%) | DNA elements (%) | Unclassified (%) | Small RNA (%) | Satellites (%) | Simple Repeats (%) | Low Complexity (%) | Total (%) | GenBank Accession | Reference |
|---|---|---|---|---|---|---|---|---|---|---|---|---|
| *Uloborus diversus* | 0.05 | 2.48 | 3.89 | 14.17 | 43.02 | 0.04 | 0.02 | 0.86 | 0.15 | 64.68 | Unpublished | Unpublished |
| *Anelosimus studiosus* | 0.71 | 1.06 | 0.38 | 7.94 | 24.06 | 0.67 | 0.1 | 0.87 | 0.28 | 35.98 | GCA_008297655.1 | Purcell and Pruitt, (2019) |
| *Araneus ventricosus* | 0.54 | 2.28 | 1.88 | 14.45 | 31.05 | 0.17 | 0.62 | 0.5 | 0.13 | 55.96 | GCA_013235015.1 | Kono, *et al*. (2019) |
| *Argiope bruennichi* | 0.08 | 1.6 | 0.76 | 6.27 | 20.52 | 0 | 0.63 | 1.58 | 0.42 | 34.64 | GCA_015342795.1 | Sheffer, *et al*. (2020) |
| *Dysdera silvatica* | 1.44 | 12.33 | 1.09 | 19.58 | 24.49 | 0.2 | 0 | 0.87 | 0.14 | 60.03 | GCA_006491805.1 | Sanchez-Herrero, *et al*. (2019) |
| *Latrodectus hesperus* | 1.75 | 2.33 | 0.27 | 7.03 | 6.83 | 0.52 | 0 | 1.61 | 0.49 | 20.97 | GCA_000697925.2 | No current papers |
| *Loxosceles reclusa* | 1.81 | 8.74 | 1.52 | 10.23 | 13.25 | 1.29 | 0.02 | 0.15 | 0.03 | 36.51 | GCA_001188405.1 | No current papers |
| *Nephila clavipes* | 0.85 | 1.14 | 0.59 | 13.71 | 18.11 | 0.48 | 0.04 | 0.65 | 0.15 | 36.61 | GCA_002102615.1 | Babb, *et al*. (2017) |
| *Parasteatoda tepidariorum* | 2.75 | 1.39 | 0.52 | 6.9 | 22.14 | 0.29 | 0.47 | 0.9 | 0.29 | 36.79 | GCA_000365465.3 | Schwager, *et al*. (2017) |
| *Pardosa pseudoannulata* | 0.8 | 1.73 | 1.71 | 16.55 | 23.16 | 1.12 | 0.24 | 2.42 | 0.44 | 48.61 | GCA_008065355.1 | Yu, *et al*. (2019) |
| *Stegodyphus dumicola* | 0.7 | 4.3 | 8.97 | 16.17 | 16.61 | 0.19 | 0.14 | 0.9 | 0.16 | 58.98 | GCA_010614865.1 | Liu, *et al*. (2019) |
| *Stegodyphus mimosarum* | 0.57 | 3.5 | 6.91 | 18.77 | 24.6 | 1.11 | 0.05 | 0.63 | 0.14 | 56.91 | GCA_000611955.2 | Sanggaard, *et al*. (2014) |
| *Trichonephila antipodiana* | 1.11 | 3.63 | 3.47 | 22.58 | 22.17 | 0.72 | 0.13 | 1.08 | 0.19 | 59.21 | | |

*Repeat statistics not reported for Acanthoscurria geniculata, Dolomedes plantarius, or Oedothorax gibbosus
